# Dynamics and plasticity of stem cells in the regenerating human colonic epithelium

**DOI:** 10.1101/2023.12.18.572103

**Authors:** Koen C. Oost, Maurice Kahnwald, Silvia Barbiero, Gustavo de Medeiros, Simon Suppinger, Veronique Kalck, Ludivine Challet Meylan, Sebastien A. Smallwood, Michael B. Stadler, Prisca Liberali

## Abstract

The human intestinal epithelium is a tissue with rapid turnover. Its complex regenerative process and differentiation trajectories have been challenging to study due to its inaccessibility and lack of temporal sampling. To this end, we developed a workflow to culture adult stem cell-derived human intestinal organoids from single cells to maturation. Extensive characterization of our model system indicated a transient regenerative response, followed by differentiation into all mature cell lineages. This switch is accompanied by a transition between two stem cell states. High-content screening and comparison to *in vivo* studies revealed that an initial fetal-like state is crucial for achieving successful regeneration, while the subsequent adult-like state is vital for maintaining a balance of cell lineages and continuous support of crypt-morphogenesis. Taken together, this study highlights the extensive plasticity of the intestinal epithelium and paves the way for further studies of human intestinal regeneration and its deregulation in pathologies.

## Introduction

The intestinal epithelium lines the inner surface of the intestine, supported by a complex mesenchyme, and plays a critical role in the absorption of nutrients and water as well as in the regulation of hormonal and immune responses. Under homeostasis, it undergoes rapid turnover every 5-7 days^1^, making it a highly dynamic tissue which requires constant proliferation, migration, and differentiation towards secretory and absorptive cell lineages. Such a dynamic nature highlights the importance of temporal resolution to understand its functions. Under injury and damage, the intestinal epithelium repairs itself by dedifferentiation of mature cells from all lineages^2,3^ to a highly proliferative regenerative state positive for Yes associated protein 1 (YAP1). YAP1 is a transcriptional coactivator, involved in regenerative processes of various tissues, that initiates widespread transcriptional changes, including upregulation of Annexin A1 (ANXA1) and Clusterin (CLU)^4–10^.

Distinct differences in cell type composition, morphology, and fundamental functions divide the intestinal tract into the small intestine and the colon^11–15^. The small intestine is characterized by finger-like protrusions called villi, which maximize the surface area for nutrient absorption through numerous absorptive enterocytes. The colon presents a contrasting morphology, with a luminal surface that lacks villi and is primarily constituted by goblet cells and colonocytes. Notably, compared to the small intestine the colon is devoid of WNT-ligand producing Paneth cells. Furthermore, a wide variety of pathologies are biased towards the colonic part of the human intestine, such as Inflammatory Bowel Disease (IBD) including Ulcerative Colitis, as well as colorectal cancer (CRC). Despite these differences, a significant portion of our current understanding of mechanistic processes comes from studies carried out in the small intestine of the mouse^11^ and it has been difficult to evaluate how conserved these processes are in the human intestine, and especially in the colon^2^.

To overcome this limitation, researchers have turned to 3-dimensional (3D) organoid systems. Organoids are models derived from induced pluripotent, fetal, or adult stem cells (iPSC, FSC, or ASC, respectively) that mimic the function and cell type composition of their *in vivo* counterparts remarkably. They enable genetic, pharmacological, or mechanical perturbations with high temporal sampling^16–19^. Human intestinal organoids have been challenging to establish, necessitating sophisticated culture conditions. While continuous advancements are made in improving culture protocols, many human organoid models suffer from an unbalanced cell type composition and failure to robustly establish correct spatial organization of cell types. Importantly, an in-depth comparison of organoids derived from these protocols to *in vivo* tissue is missing, leading to unclear translatability of obtained results.

To improve ASC-derived human intestinal organoids, we focused on signaling pathways that improve the regenerative capacity of the intestinal epithelium and the balanced differentiation towards the major cell lineages. Recent studies have highlighted context-dependent significance of the epidermal growth factor (EGF)-like ligands EGF and neuregulin 1 (NRG1) in the intestinal epithelium^20–22^. Especially NRG1, an ERBB3/4 ligand, has been shown to promote regeneration after damage in the mouse small intestine as well as elevated maturation and diversity of cells in human (fetal) small intestinal organoids^20–22^.

We thus applied high-content screening to probe the effects of activation and inhibition of ERBB-receptors at defined time windows. We found that an initial signal, likely through EGF-induced EGFR homodimers, is essential for survival and a subsequent switch to elevated ERBB3-signaling during homeostasis improves stem cell maintenance and secretory-lineage differentiation. Building upon this insight, we formulated a medium which enables continuous cultures from single cells into mature homeostatic organoids with high similarity to *in vivo* tissue. In-depth analysis of their maturation trajectory revealed a transition from a regenerative to a homeostatic phase after dedifferentiation from all mature lineages, enabling insights into previously inaccessible dynamic processes of the human intestinal epithelium. Our work showcases that this switch is connected to two distinct stem cell states. An initial state resembling signatures of fetal stem cells is crucial for achieving successful regeneration, while a subsequent state akin to adult stem cells is vital for maintaining a proper balance of cell lineages and crypt-morphogenesis. This highlights the complex tissue scale dynamics a regenerative epithelium undergoes to reach a homeostatic multicellular system after damage. Furthermore, the developed workflow provides a valuable patient-derived model system to assess disruptions of this process during disease modelling, precision diagnostics, and pre- clinical safety screening.

## Results

### Development of a niche-inspired medium through dynamic EGF-like ligand supplementation

To improve not only the cell type composition but also tissue morphogenesis and spatial organization of the cells, we first investigated the effects of NRG1 (isoform 1) supplementation on human colonic organoids in medium with or without EGF, analyzed with high-content microscopy (Figure 1A, light pink box). To quantify organoid phenotypes and crypt- morphogenesis with more sensitivity, we developed an image analysis workflow to extract organoid-level cell-type marker expression levels and morphological features including the number of crypts, as well as their total length from more than 40,000 organoids used in this study (Figure S1A, Methods). In intestinal organoids, a crypt-like structure usually represents a favorable phenotype, as it indicates an actively proliferating stem cell (SC) compartment resembling the SC-niche of *in vivo* tissue.

**Figure 1.**
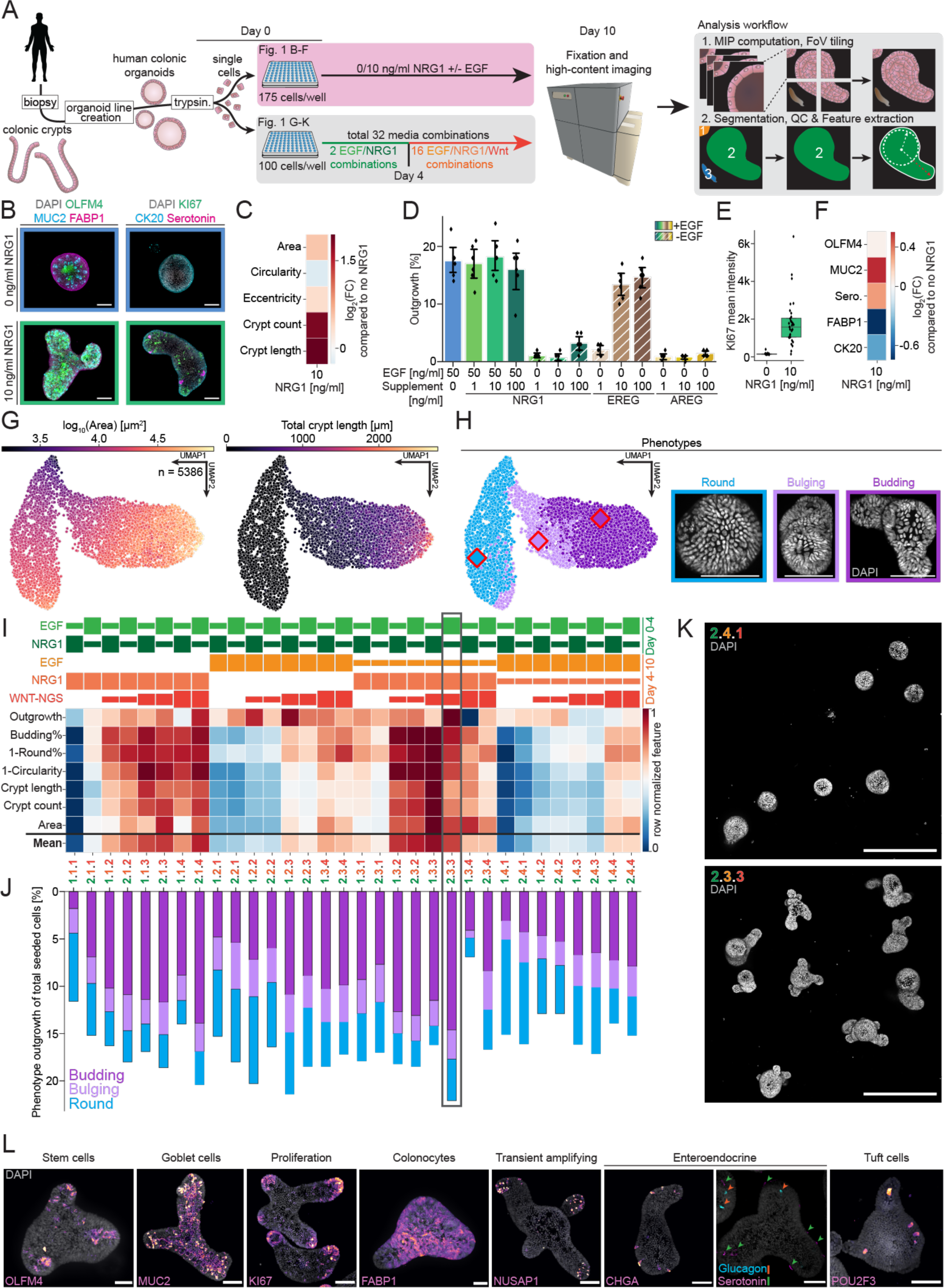
Development of a niche-inspired medium composition through dynamic EGF-like ligand supplementation. (**A**) Schematic of the experimental workflow to test effect of NRG1, EGF and WNT-NGS on human colonic organoid maturation from single cells. (**B-F**) Results of constant NRG1 supplementation. (**B**) Representative images of 10-day old human colonic organoids grown from single cells without (upper row) and with (lower row) NRG1 supplementation. Scale bars, 100 µm. (**C**) Log2(fold change) of morphological features of organoids grown with NRG1 supplementation compared to without. (**D**) Outgrowth efficiency of organoids from single cells under constant EGF-like ligand influence. Bars with stripes denote conditions with 50 ng/ml EGF supplementation. Individual data points present outgrowth in one well. Data denotes mean and 95% confidence interval. (**E**) KI67 mean intensity of organoids matured without or with NRG1. Bar denotes median ± quartiles. (**F**) Log2(fold change) of marker expression of organoids grown with NRG1 supplementation compared to without. (**G- K**) Results of time-dependent EGF-like ligand supplementation on human colonic organoids. (**G**) Selected features highlighted on a Uniform Manifold Approximation and Projection (UMAP) based on the shape-feature space of mature colonic organoids grown in 32 different conditions (see Table 1). (**H**) Three major observed phenotypes (‘Round’: light blue, ‘Bulging’: light pink, ‘Budding’: purple) derived from shape-features and exemplary images of colonic organoids within this phenotype. Diamonds with red border show coordinates of the organoids used as phenotype examples. Scale bars, 100 µm. (**I**) Heatmap of row-normalized features associated with successful crypt-morphogenesis among all 32 tested conditions (see Table 1). Size of bars above heatmap represents the concentrations of EGF and NRG1 until day 4 (green), as well as EGF, NRG1 and WNT-NGS from day 4 to day 10 (orange). (**J**) Outgrowth efficiency of tested conditions color-coded by the phenotype of surviving organoids. Grey rectangle indicates medium combination 2.3.3 (see Table 3), which is used for further experimentation. (**K**) Representative whole-well images of a poor (2.4.1) and well (2.3.3) performing medium combination. Scale bars, 500 µm. (**L**) Representative images of organoids grown in medium combination 2.3.3 immunostained for proliferation and major human colonic cell types. Rare cells positive for Glucagon or Serotonin are indicated with orange and green arrows, respectively. Scale bars, 50 µm. All images are maximum intensity projections (MIPs) of z-stacks acquired on a spinning-disk confocal system.

**Table 1:**
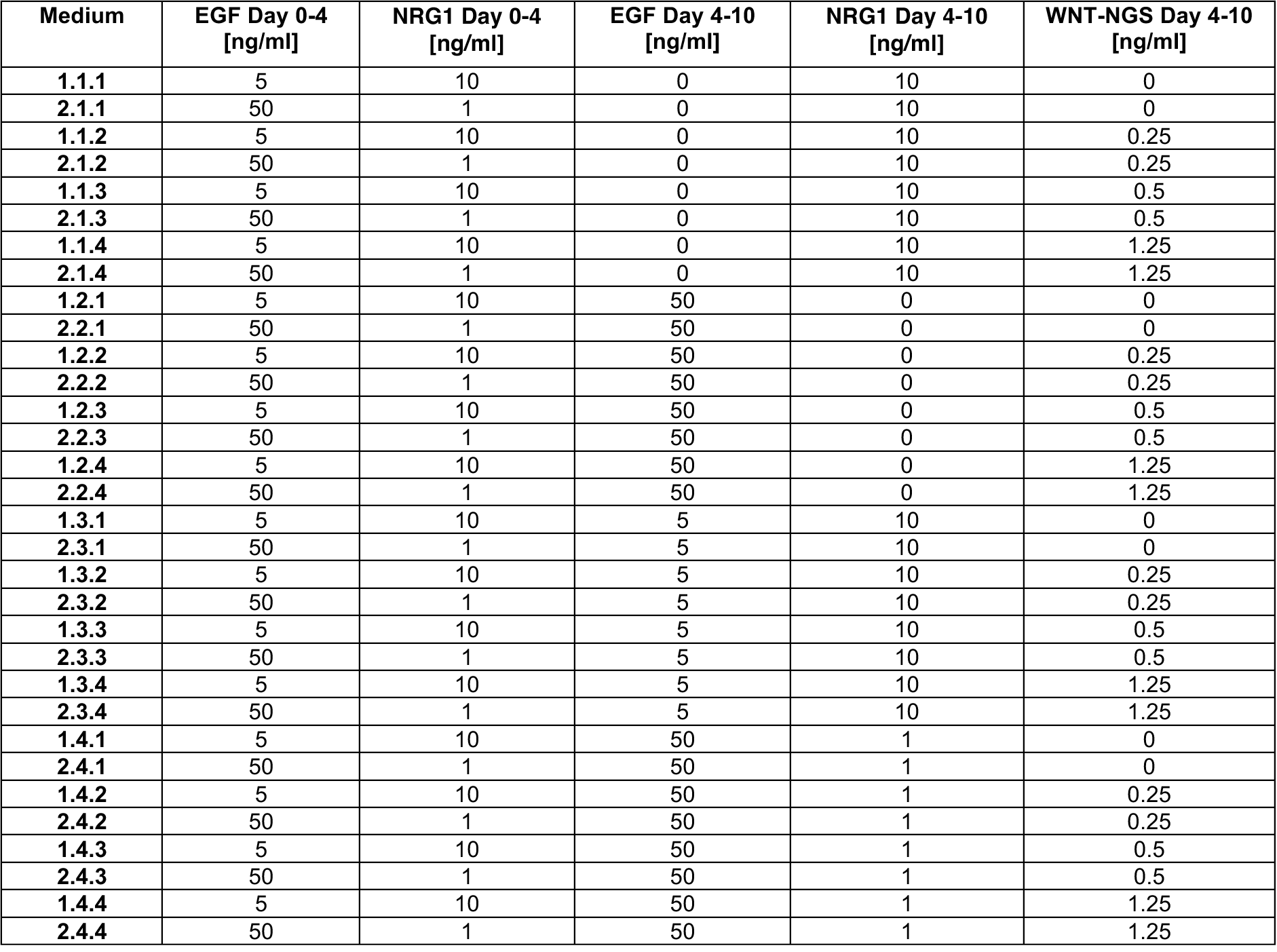
Human colon organoid media test. Tested media combinations and their concentrations of NRG1, EGF, and WNT-NGS in addition to the base medium (Methods) for human colon organoids. Medium combination 2.3.3 has been used for further work and was termed Start (Day 0-4 medium) and Balance (Day 4-10 medium).

| Medium | EGF Day 0-4 [ng/ml] | NRG1 Day 0-4 [ng/ml] | EGF Day 4-10 [ng/ml] | NRG1 Day 4-10 [ng/ml] | WNT-NGS Day 4-10 [ng/ml] |
| --- | --- | --- | --- | --- | --- |
| 1.1.1 | 5 | 10 | 0 | 10 | 0 |
| 2.1.1 | 50 | 1 | 0 | 10 | 0 |
| 1.1.2 | 5 | 10 | 0 | 10 | 0.25 |
| 2.1.2 | 50 | 1 | 0 | 10 | 0.25 |
| 1.1.3 | 5 | 10 | 0 | 10 | 0.5 |
| 2.1.3 | 50 | 1 | 0 | 10 | 0.5 |
| 1.1.4 | 5 | 10 | 0 | 10 | 1.25 |
| 2.1.4 | 50 | 1 | 0 | 10 | 1.25 |
| 1.2.1 | 5 | 10 | 50 | 0 | 0 |
| 2.2.1 | 50 | 1 | 50 | 0 | 0 |
| 1.2.2 | 5 | 10 | 50 | 0 | 0.25 |
| 2.2.2 | 50 | 1 | 50 | 0 | 0.25 |
| 1.2.3 | 5 | 10 | 50 | 0 | 0.5 |
| 2.2.3 | 50 | 1 | 50 | 0 | 0.5 |
| 1.2.4 | 5 | 10 | 50 | 0 | 1.25 |
| 2.2.4 | 50 | 1 | 50 | 0 | 1.25 |
| 1.3.1 | 5 | 10 | 5 | 10 | 0 |
| 2.3.1 | 50 | 1 | 5 | 10 | 0 |
| 1.3.2 | 5 | 10 | 5 | 10 | 0.25 |
| 2.3.2 | 50 | 1 | 5 | 10 | 0.25 |
| 1.3.3 | 5 | 10 | 5 | 10 | 0.5 |
| 2.3.3 | 50 | 1 | 5 | 10 | 0.5 |
| 1.3.4 | 5 | 10 | 5 | 10 | 1.25 |
| 2.3.4 | 50 | 1 | 5 | 10 | 1.25 |
| 1.4.1 | 5 | 10 | 50 | 1 | 0 |
| 2.4.1 | 50 | 1 | 50 | 1 | 0 |
| 1.4.2 | 5 | 10 | 50 | 1 | 0.25 |
| 2.4.2 | 50 | 1 | 50 | 1 | 0.25 |
| 1.4.3 | 5 | 10 | 50 | 1 | 0.5 |
| 2.4.3 | 50 | 1 | 50 | 1 | 0.5 |
| 1.4.4 | 5 | 10 | 50 | 1 | 1.25 |
| 2.4.4 | 50 | 1 | 50 | 1 | 1.25 |

Addition of NRG1 to medium containing 50 ng/ml EGF resulted in larger colonic organoids with higher crypt count and crypt length (Figures 1B, 1C, and S1B). The outgrowth efficiency (∼17%) in our base-medium was not affected by NRG1, and its supplementation alone did not rescue outgrowth failure due to absent EGF (Figure 1D). However, the high-affinity EGFR-ligand EREG induced organoid formation in comparison to no rescue with the lower-affinity ligand AREG within the tested concentration range (Figure 1D)^25^. Staining for well-established markers of proliferation (KI67), SCs (OLFM4), secretory cells (MUC2/Serotonin) and absorptive cells (FABP1/CK20) in media containing NRG1 revealed a substantial increase in KI67 expression (∼25 fold), and a bias towards the secretory lineage with dampened absorptive-marker expression compared to media containing no NRG1 (Figures 1B, 1E, 1F, and Figure S1B). Taken together, these results indicate that while EGFR-activation is necessary for initial organoid outgrowth, NRG1 supplementation increases differentiation towards secretory cell lineages and maintains a proliferative state in crypt-regions of organoids.

To further finetune EGF and NRG1 concentrations, we varied their ratios in two different time windows: (i) directly after single cell seeding until day 4 to improve outgrowth from single cells and (ii) from day 4 to fixation at day 10 to support crypt-morphogenesis. Furthermore, while we identified that exogenous WNT-pathway activation is necessary during the initial time window (data not shown), we tested four varying concentrations of WNT-NGS after day 4 (Table 1)^26^. Taking into consideration all aforementioned conditions, we performed an imaging-based screen with a total of 32 tested media (Figure 1A (light gray box), Figures 1G- 1K, Table 1). The UMAP embedding of image-based morphological features from 5386 organoids after quality control (Methods) and visual assessment of the samples indicated three main phenotypic clusters – ‘Round’, ‘Bulging’, ‘Budding’ (Figures 1G and 1H, Methods). ‘Round’ organoids failed to develop crypt-like structures, while ‘Bulging’ organoids show first indications of crypt-morphogenesis due to decreased circularity and precede ‘Budding’ organoids^8^. Organoids in the ‘Budding’ phenotype developed at least one crypt-like structure. All media combinations were analyzed based on outgrowth efficiency, relative percentage of organoids belonging to the ‘Budding’ cluster, as well as crypt parameters among others (Figure 1I).

We found that WNT-NGS supplementation during the second phase (4-10 days) of organoid maturation is necessary for increased survival and crypt-morphogenesis. This is consistent with *in vivo* studies showing that colonic epithelial cells rely predominantly on mesenchymal WNT, rather than secreting WNT-ligands themselves during homeostasis^18,27–31^.

Varying EGF to NRG1 ratios until day 4 has low impact on the final phenotype. A shift occurs after day 4, where lower EGF and higher NRG1 concentrations promote crypt-morphogenesis. Based on these observations and the absolute number of budding organoids per well (Figure 1J), medium combination 2.3.3 emerged as the most promising candidate (Figure 1K). The staining of cell type markers in this medium shows the presence of all major cell-types in their expected location and abundance (Figure 1L)^14,15^. Henceforth, we named the initial medium *Start*, followed by the *Balance* medium from day 4 onwards (Table 3). To examine the wide applicability of the *Start*/*Balance* medium, we utilized it on a total of 47 biopsy-derived organoid lines obtained from normal intestinal epithelial tissue across all sublocations of the colon and successfully maintained continuous cultures for all patient-derived organoid lines (Figure S1C, Methods). Comparing the workflow of this study to other established protocols and a commercial option showed superior outgrowth and a more pronounced crypt-morphology in the *Start/Balance* regime after outgrowth from single cells in the context of this experiment (Figure S1D)^18,19^.

To have real time understanding of the organoid development, we created a H2B-iRFP-670 human colon organoid reporter line for nuclei (Methods). Real-time imaging of 44 organoids on a light sheet microscope over 200 hours of maturation from single cells revealed sub- organoid dynamics (Videos S1 and S2). We found that 8 (18.2%) out of 44 organoids did not survive the long-term imaging even though starting from viable cells. In the other 36 organoids, initially, cells exhibited high mobility and proliferated throughout the sphere indiscriminately. From day 5 onwards, crypt-morphogenesis emerged, leading to increased proliferation in this region. Additionally, we noted collective cell behaviors, including dynamic rotation of cells above the crypt neck, contrasting less mobility in the crypt. These observations underscore the complex and dynamic nature of organoid maturation.

To establish a comparable medium regime for human ASC-derived small intestinal organoids, we revisited a similar strategy as for colonic organoids (Figures S1E-S1I, Table 2). However, contrary to a recently described protocol^33^, duodenal small intestinal organoids are not dependent on WNT-NGS after initial outgrowth, and we thus removed WNT-NGS from the balance medium. For this reason, we have tested 8 media combinations for their ability to induce crypt-morphogenesis. Notably, the top-performing medium combination was identical to the one identified for colonic organoids, but devoid of WNT-NGS in the *Balance* medium and instead supplemented with all-trans retinoic acid (atRA) to increase enterocyte differentiation^5^ (Figure S1H). Furthermore, we find all major cell types, including LYZ1^+^ Paneth cells, in mature small intestinal organoids (Figure S1I). Finally, we found that applying the medium regime to colonic or small intestinal organoids of *Rattus rattus* (Figures S1J and S1K) and colonic *Mus musculus* organoids (Figures S1L and S1M) led to a stable outgrowth into organoids with all major cell types. Taken together, the developed workflow allows the quantitative analysis of human intestinal organoid development from single cells towards fully mature organoids in a scalable and robust manner.

**Table 2:** Human small intestine organoid media test. Tested media combinations and their concentrations of NRG1, EGF, and WNT-NGS in addition to the base medium (Methods) for human small intestinal organoids. Medium combination 2.3 has been used for further work and was termed Start (Day 0-4 medium) and Balance (Day 4-10 medium).

| Medium | EGF Day 0-4 [ng/ml] | NRG1 Day 0-4 [ng/ml] | EGF Day 4-10 [ng/ml] | NRG1 Day 4-10 [ng/ml] | WNT-NGS Day 4-10 [ng/ml] |
| --- | --- | --- | --- | --- | --- |
| 1.1 | 5 | 10 | 0 | 10 | 0 |
| 2.1 | 50 | 1 | 0 | 10 | 0 |
| 1.2 | 5 | 10 | 50 | 0 | 0 |
| 2.2 | 50 | 1 | 50 | 0 | 0 |
| 1.3 | 5 | 10 | 5 | 10 | 0 |
| 2.3 | 50 | 1 | 5 | 10 | 0 |
| 1.4 | 5 | 10 | 50 | 1 | 0 |
| 2.4 | 50 | 1 | 50 | 1 | 0 |

**Table 3:** Final human intestinal medium composition. Composition of the final medium from day 0 to day 4 (termed *Start*) and from day 4 to day 10 (termed *Balance)* for the colonic and small intestinal organoids. Ingredients should be diluted with DMEM/F12 (+15 mM HEPES). Derived from combination 2.3.3 from Table 1 and 2.3 from Table 2, respectively. *Human R-Spondin1 was a kind gift of Novartis. Recombinant human R-Spondin3 (250 ng/ml *Start* and 125 ng/ml *Balance*) works but was not extensively tested in long-term cultures. **murine EGF (PeproTech, 315-09-100ug) can also be used.

| Compound | *Start (day 0 to day 4)* |  | *Balance (day 4 to day 10)* |  |
| --- | --- | --- | --- | --- |
|  | Colon | Small intestine | Colon | Small intestine |
| Next-Generation Surrogate WNT (WNT-NGS) | 0.5 nM | 0.5 nM | 0.1 nM | - |
| human R-Spondin1* | 1 µg/ml | 1 µg/ml | 0.5 µg/ml | 0.5 µg/ml |
| murine Noggin | 100 ng/ml | 100 ng/ml | 100 ng/ml | 100 ng/ml |
| N-Acetylcysteine (NAC) | 1.25 mM | 1.25 mM | 1.25 mM | 1.25 mM |
| A83-01 | 0.5 µM | 0.5 µM | - | - |
| human IGF1 | 100 ng/ml | 100 ng/ml | 100 ng/ml | 100 ng/ml |
| human FGF2 | 50 ng/ml | 50 ng/ml | 50 ng/ml | 50 ng/ml |
| B27 supplement | 1x | 1x | 1x | 1x |
| human Gastrin | 25 nM | 25 nM | 25 nM | 25 nM |
| human EGF** | 50 ng/ml | 50 ng/ml | 5 ng/ml | 5 ng/ml |
| human NRG1 | 1 ng/ml | 1 ng/ml | 10 ng/ml | 10 ng/ml |
| Y-27632 | 10 µM | 10 µM | - | - |
| all-trans retinoic acid (atRA) | - | - | - | 2.5 µM |
| Penicillin/Streptomycin | 100 U/ml | 100 U/ml | 100 U/ml | 100 U/ml |
| GlutaMAX supplement | 1x | 1x | 1x | 1x |

### Image-based pseudotime analysis indicates two distinct maturation trajectories

To characterize colonic organoid maturation dynamics, we performed multiple high-content imaging time courses over a period of ten days with 48-hour sampling intervals (Figure 2A). To avoid averaging features by fixation timepoints across asynchronous maturation stages we used trajectory inference algorithms to order organoids based on pseudotime and infer their maturation trajectories (Figures 2A, 2B and S2A-S2C, Methods)^34^. Through this workflow, we identified two distinct maturation trajectories, resulting in the formation of either round organoids (‘Round’, yellow trajectory) or organoids which initially start to bulge and eventually undergo crypt-morphogenesis (‘Budding’, dark blue trajectory). Of note, the removal of crypt-associated features (Figure S1A, Methods), such as the number of crypts or crypt length, results in the total inability to detect the bifurcation event (Figures S2D and S2E). Following morphological features over pseudotime revealed that both trajectories, similar to mouse small intestinal organoids^5,32^, show initially a round cyst-like morphology (Figure 2E). Morphological symmetry is broken at pseudotime bin 4/5 (∼5-6 days in estimated real-time) exclusively in the ‘Budding’ trajectory at an average diameter of 85 µm. Shortly thereafter at pseudotime bin 6, the first crypts are detected in this branch. The crypts then continue to elongate until the experimental end point, indicating an active SC population. ‘Round’ organoids, however, do not undergo crypt-morphogenesis but continuously grow throughout the experimental timeframe although at a lower rate.

**Figure 2.**
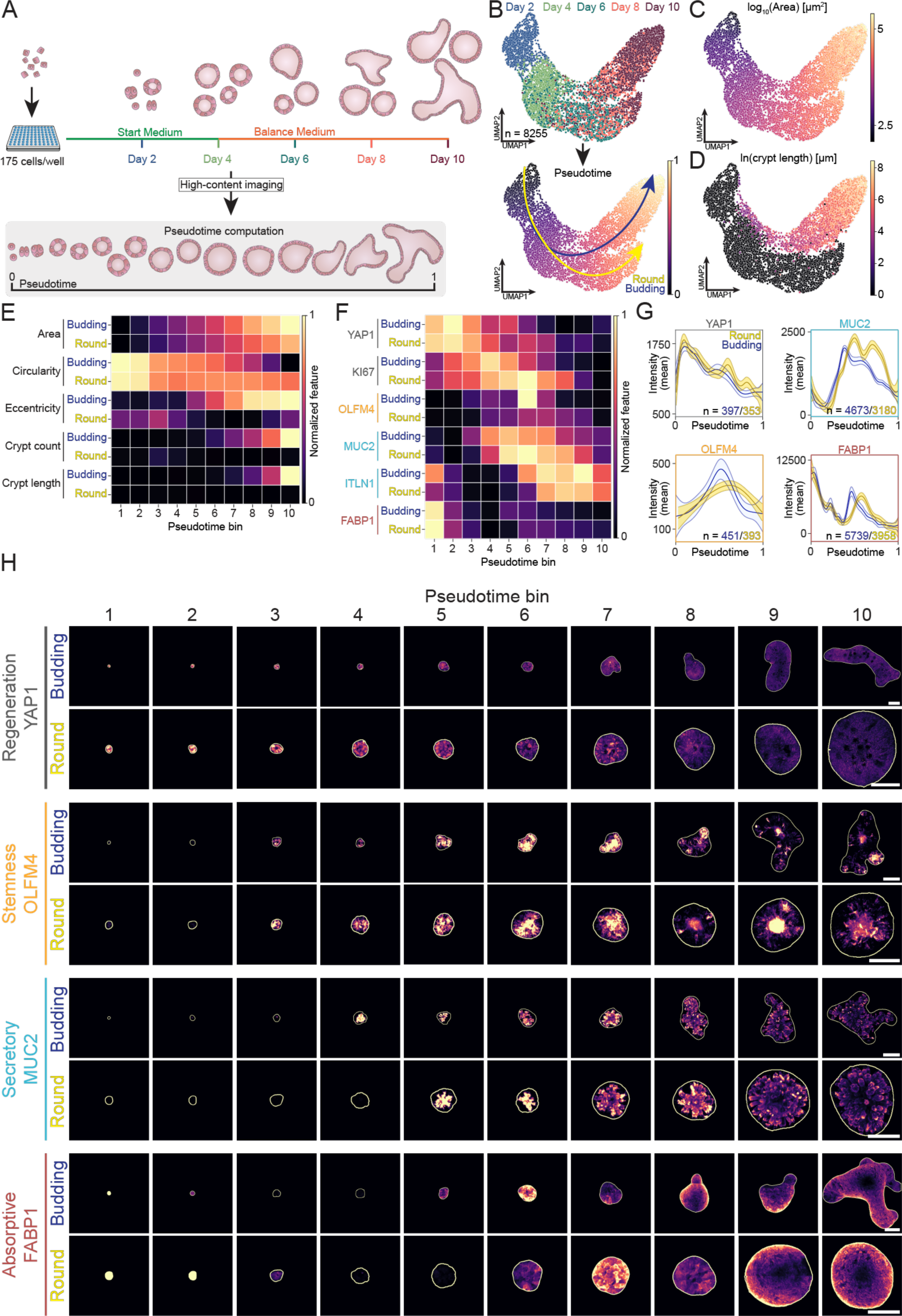
Image-based pseudotime analysis indicates two distinct maturation trajectories. (**A**) Schematic representation of the imaging time course setup from single cells. After 4 days in *Start* and 6 days in *Balance* medium (see Table 3), colonic organoids were fixed at day 10 with subsequent high-content imaging and pseudotime analysis. (**B**) Organoids from time course experiments placed in an UMAP based on their shape- feature space color-coded by the fixation timepoint (upper panel) and their resulting pseudotime (lower panel). Slingshot indicated two maturation trajectories leading to either ‘Round’ (yellow) or ‘Budding’ (blue) organoids. (**C- D**) Selected features highlighted on a UMAP based on the shape-feature space. (**E-F**) Normalized morphological features (D) and organoid-level mean intensities of markers associated with regeneration, proliferation, and cell types over pseudotime bins separated by both trajectories. (**G**) Predicted mean intensity by a generalized additive model (GAM) including 95% confidence intervals of markers associated with regeneration and cell type identity over pseudotime. (**H**) Representative images of marker expression in organoids along pseudotime bins in both trajectories. White outline represents the segmentation mask. Images are maximum intensity projections (MIPs) of z-stacks acquired on a spinning-disk confocal system. Scale bars, 100 µm.

To identify the various phases of the maturation trajectories and cell type emergence, we used immunofluorescence analysis of markers associated with active damage-induced regeneration (YAP1)^7^, proliferation (KI67), and major cell types of the human intestine (SC: OLFM4; goblet cells (GC): MUC2/ITLN1; enteroendocrine cells (EEC): CHGA; colonocytes (C): FABP1) (Figures 2F-2H, S2F and S2G). As our framework allows the monitoring of multiple markers along common maturation trajectories, it revealed a high regenerative potential in organoids prior to crypt-morphogenesis, characterized by nuclear localization of YAP1 (Figure S2H) and enhanced proliferation marked by KI67 (Figures 2F-2H, S2F and S2G). The inactivation and translocation of YAP1 out of the nucleus coincides with the expression of the first canonical cell type markers, such as for SC (OLFM4^+^) and GC (MUC2^+^) and hints towards an initial YAP1^+^ regenerative phase, void of canonical cell type markers. As the maturation progresses, proliferation becomes restricted to the crypt regions, reflecting the distribution of intestinal SC (S2G). At the cell type level, OLFM4^+^ cells emerge at a similar time as MUC2^+^ positive secretory cells with minor differences between both trajectories (Figures 2F-2H and S2F, S2G). Shortly after, first absorptive cells (FABP1^+^), EECs (CHGA^+^) and mature GCs (ITLN1^+^) are detected (S2G). In contrast to the small intestine, the colon has GCs distributed throughout the epithelial lining^35^, whereas colonocytes (FABP1^+^) preferentially localize away from the crypts. This spatial organization is conserved in ‘Budding’ organoids, however, not in ‘Round’ organoids. This may indicate that, while organoids are able to establish all major cell types in both trajectories, an expected cell type-gradient is not developed in the ‘Round’ branch. These findings provide unprecedented insights into the spatiotemporal dynamics of human colon organoid maturation from single cells. Moreover, we found that increased expression and nuclear localization of YAP1 indicates a transient regenerative state after seeding of single cells before the emergence of the first canonical cell types.

### Differential ERBB-receptor activation influences lineage specification in human colonic organoids

We and others report vast effects of EGF-like ligands on model systems of the intestinal epithelium (Figure 1, reviewed by Abud and colleagues^24^). These ligands bind to receptors of the ERBB-family and induce their homodimerization, their heterodimerization with ERBB2, as well as oligomers of higher order which signal into a complex downstream network including PI3K, MAPK, and JAK-STAT signaling (Figure S3A)^23,24,36^. However, a detailed understanding of their effects on the SC-compartment and the lineage specification in the human intestinal epithelium remains elusive. To this end, we took advantage of the pharmacological accessibility and scalability of our workflow and treated human colonic organoids with antibodies inhibiting EGFR (Cetuximab), ERBB2 (Pertuzumab), and ERBB3 (Lumretuzumab) after seeding of single cells until their maturation at day 10 (Figure 3A, Methods). The treatments have been conducted either separately, or in a combination to inhibit reported heterodimers (i.e., Cetuximab + Pertuzumab against the EGFR/ERBB2 heterodimer or Pertuzumab + Lumretuzumab against the ERBB2/3 heterodimer). Of note, we could not detect expression of the other receptor for NRG1, ERBB4, by qPCR and single cell RNA-sequencing in mature organoids, in accordance with previous reports (data not shown)^20,37^.

**Figure 3.**
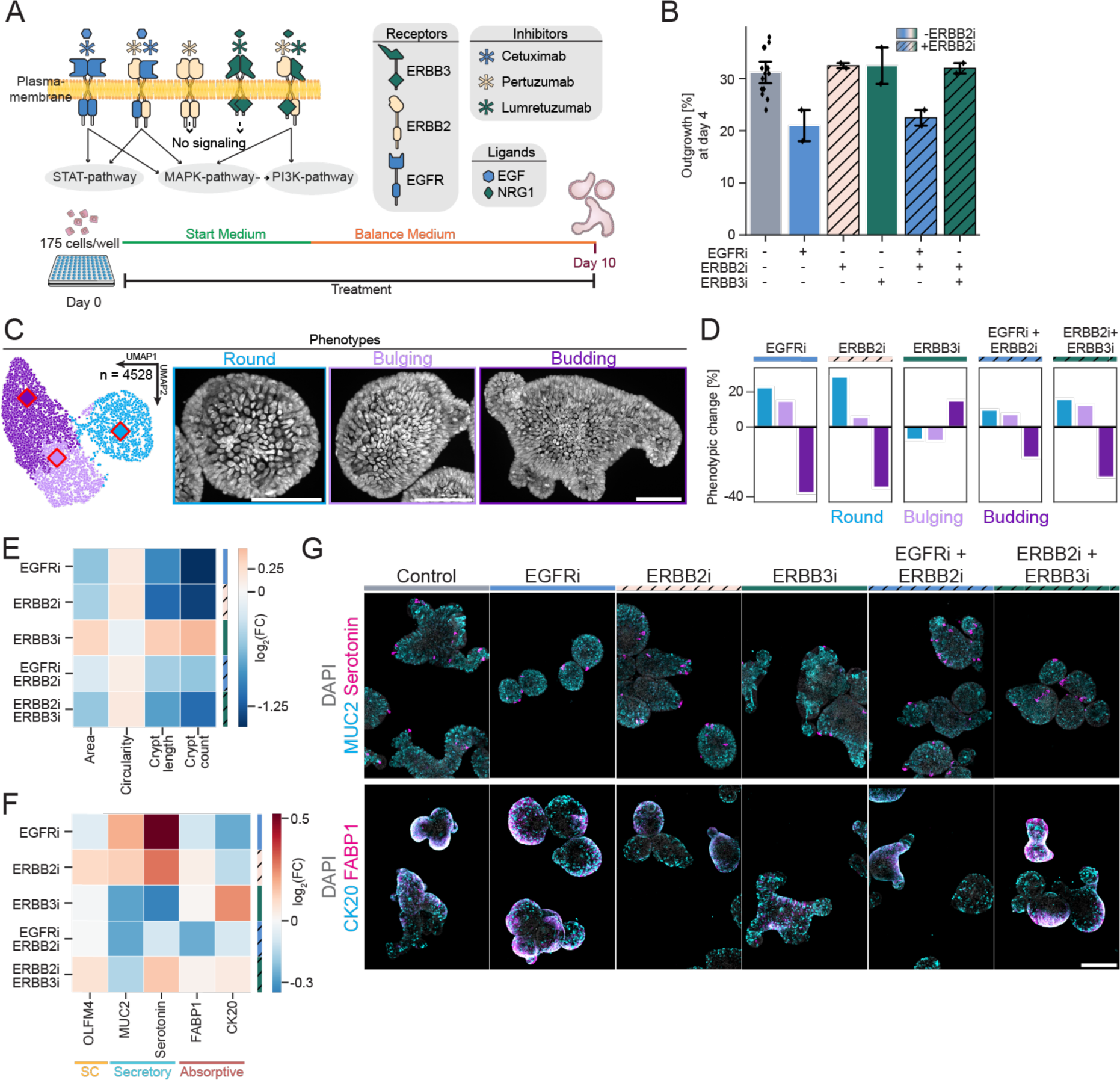
Differential ERBB-receptor activation influences lineage specification in human colonic organoids. (**A**) Simplified, schematic representation of the ERBB-receptor class signaling in the intestinal epithelium and the experimental approach. Single cells derived from mature organoids were seeded at day 0 and grown for 10 days. During this time, ERBB receptors (EGFR, ERRB2, and ERBB3) were inhibited by antibody treatments either as a single inhibition, or in combination with ERBB2 inhibitor to prevent heterodimer formation. (B) Outgrowth of the ERBB-inhibition experiment quantified at day 4 with brightfield imaging. Striped bars indicate conditions with added ERBB2 inhibition to prevent heterodimer formation. Data denotes mean and the 95% confidence interval. (**C**) Organoids from the ERBB-receptor inhibition (pooling all treatment windows and concentrations) placed in an UMAP based on their shape-feature space color-coded by derived phenotypes (‘Round’: light blue, ‘Bulging’: light pink, ‘Budding’: purple). Diamonds with red border indicate the position of examples. Scale bars, 50 µm. Related to Figure S3B. (**D**) Change in phenotypic composition of ERBB-inhibition experiments (columns) relative to the corresponding control. The color of the bar indicates the phenotype (‘Round’: light blue, ‘Bulging’: light pink, ‘Budding’: purple). (**E-F**) Heatmap showing the log2(fold change) of organoid-level morphological (**E**) and marker expression (mean intensity, **F**) features of used ERBB-receptor inhibition treatments compared to the corresponding control. (**G**) Representative images of organoids treated for the full growth-duration (day 0 to day 10) compared to the control (left column). Exemplary staining of secretory markers (top) and absorptive markers (bottom). Scale bar, 200 µm. All images are maximum intensity projections (MIPs) of z-stacks acquired on a spinning-disk confocal system.

To probe influences on the initial survival during the YAP1^+^ regenerative phase, we analyzed the outgrowth efficiency on day 4 after seeding of single cells. Most applied inhibitors did not have a negative influence on the survival during this phase, except for inhibition of EGFR (Figure 3B). This resembles our previous observation indicating that EGFR-activation is necessary for initial survival after induction of damage (Figure 1D). As additional ERBB2 inhibition did not augment this effect, our results suggest that the initial survival signal is linked to an EGFR-homodimer instead of an EGFR-ERBB2 heterodimer (Figure 3B).

At day 10, all conditions were fixed and stained for markers of SCs (OLFM4), the secretory lineage (MUC2, Serotonin), and the absorptive lineage (CK20, FABP1). Following feature extraction (Figure S3B), we identified 3 major end-point phenotypes (Figure 3C), similar to our results of the EGF-like ligand titration (Figures 1G-1K). Most treatments shifted the distribution of these phenotypes considerably compared to the control (Figure 3F). While inhibition of EGFR and ERBB2 led to a shift towards rounder organoids, ERBB3 inhibition showcased a slight shift towards organoids in the ‘Budding’ phenotype. This shift is well captured when comparing morphological features between these conditions (Figure 3E, left panel) and is accompanied by a bias towards the secretory lineage (EGFR and ERBB2 inhibition) compared to a downregulation in this lineage during ERBB3 inhibition (Figures 3E (right panel) and 3F) in accordance with our results during the supplementation of EGF-like ligands (Figure 1F).

To elucidate the effects of these inhibitions during the two major phases in our organoid maturation trajectory, regenerative and homeostatic, we inhibited ERBB-receptors during these two time-windows (treatment from day 0 to 4, or day 4 to 10, respectively). We observed that inhibition of ERBB2 largely shows effects during the regenerative phase, with opposing results of EGFR inhibition (Figures S3C and S3D). These differential time windows hint once more towards preferential homodimerization of EGFR in the human colon, linking the effects of the ERBB2 inhibition to a potential heterodimer with ERBB3. However, inhibition of ERBB3 alone did not phenocopy the ERBB2 inhibition and led only to a minor shift towards ‘Budding’ organoids (Figures S3C and S3D, left panel) with bias towards the absorptive lineage in both treatment windows (Figure S3D, right panel). These findings align with observations in conditional ERBB3 knockout mice, which demonstrated a more absorptive and hyperproliferative phenotype in the colon following a brief DSS-treatment^38^.

In summary, these findings provide insights into the complex and context-dependent importance of ERBB-receptor mediated signaling in shaping phenotypes. In combination with results from the EGF-like ligand experiments, our results show the dual importance of the EGFR homodimer, once during initial survival, and subsequently during the homeostatic phase with a likely role in crypt-morphogenesis and differentiation into the absorptive lineage. On the other hand, inhibition of ERBB3 signaling on the receptor level led to only minor changes on the phenotype compared to removal of its ligand NRG1, as well as its heterodimerization partner ERBB2, hinting at backup mechanisms to rescue ERBB3-signaling loss. Finally, these results highlight the potential of this workflow for evaluating efficacy and safety of therapeutic interventions in a non-pathological setting.

### scRNA-seq reveals all major cell lineages in the mature organoid

While preceding results of this study related supplementation of EGF and NRG1 to functions during survival and cell lineage specification, a detailed temporal understanding of the molecular events which govern the initial regenerative capacity and define lineage decision- making is lacking. To this end, we conducted a time-resolved single-cell RNA sequencing (scRNA-seq) experiment. Beginning with single cells sorted from fully mature organoids, we sequenced a total of 53640 cells after quality control at 9 time points over 10 days (Figures 4A and S4A-S4C, Methods). We performed cell-cycle correction and utilized geometric sketching to refine the UMAP visualization towards underrepresented cell types^42^ (Figures 4B and S4D-SK, Methods). We observed a clear change at the transcriptome-level when the medium is changed from *Start* to *Balance*, with minimal overlap between cells coming from either medium (Figure 4C).

**Figure 4.**
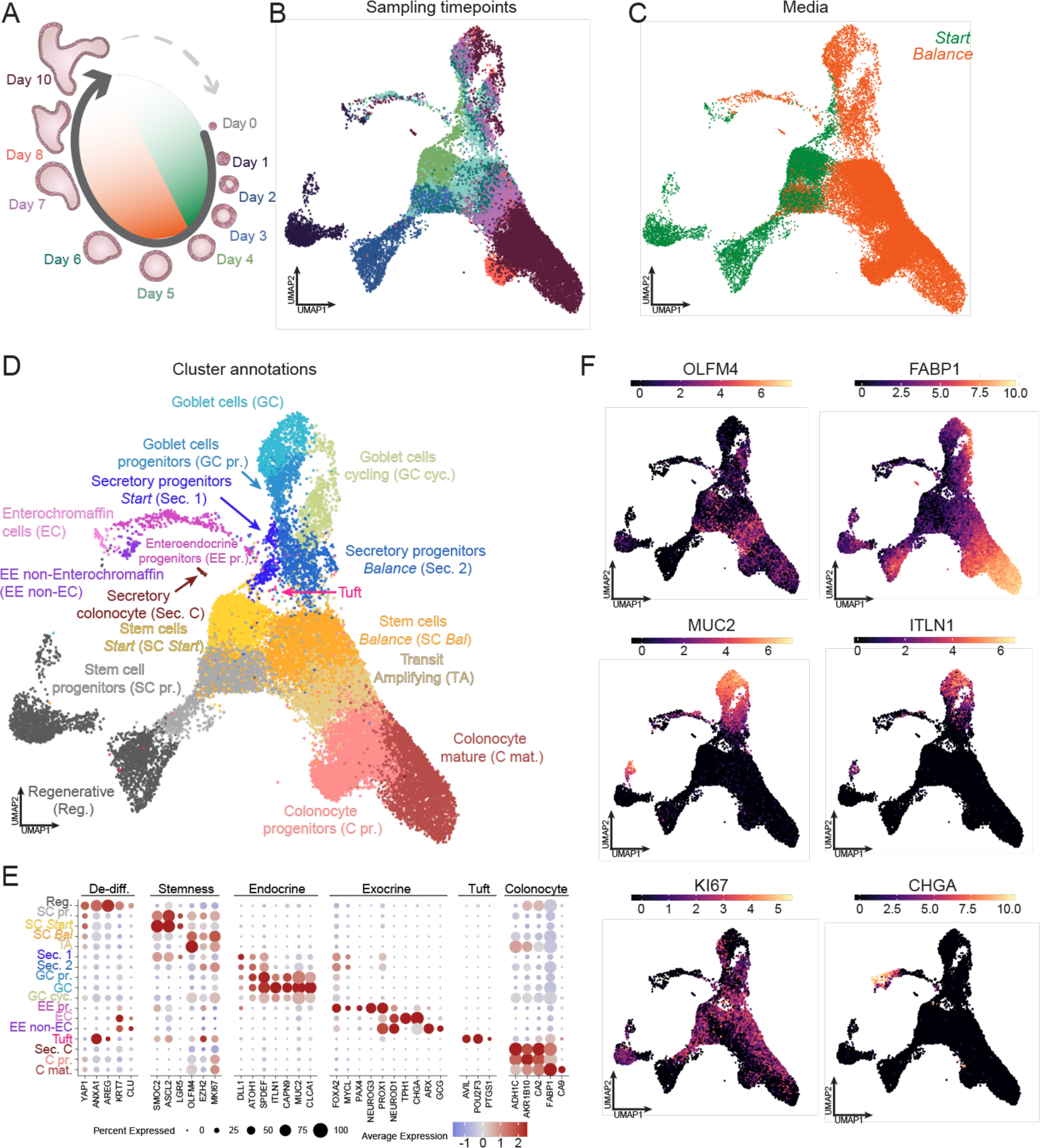
scRNA-seq reveals all major cell lineages in the mature organoid. (**A**) Schematic of the experimental layout. Mature colonic organoids were trypsinized to single cells before seeding into culture plates. Samples were harvested for single-cell RNA sequencing (scRNA-seq) on day 1, 2, 3, 4, 5, 6, 7, 8, and 10. Green inner circle represents timeframe in *Start* medium, orange time in *Balance* medium. (**B-D**) UMAP of the transcriptomic landscape of n = 53640 cells after cell-cycle correction and sketching (see Methods) color-coded by their sampling timepoint (**B**), the used medium at the timepoint of sampling (**C**), and cluster annotation (**D**). (**E**) Dot-plot of genes associated with dedifferentiation, stemness, endo/exocrine cells, as well as tuft cells and colonocytes across annotated clusters from the scRNAseq time course. Dot-color shows the average expression level while its size indicates the fraction of positive cells. (**F**) UMAP of the transcriptomic landscape after cell-cycle correction and sketching (see Methods) color-coded by major markers used in this study.

17 distinct clusters were identified and annotated to mature cell types, their progenitors, stem cells, as well as regenerative cells (Figures 4D-4F). This approach identified all major cell types, excluding a unique cluster for a recently reported BEST4^+^ population^43^. Of note we detected only a small percentage (4.19%) of GC (including all clusters that express MUC2) in the scRNA-seq compared to our imaging data (12.81% MUC2^+^ cells, Figures S4L and S4M). This likely represents a technical artefact that is introduced during singularization^45^ of organoids to single cells as a pre-processing step for scRNA-seq due to their large size or mucin-associated adhesiveness of GCs. To reinforce this observation, we report a consistent reduction in GC proportions when assessing MUC2^+^ cells through Fluorescence Activated Cell Sorting (FACS) data immediately after singularization (Figure S4M).

Nonetheless, our analysis yielded unique insights into the maturation process of various cell types. Shortly after seeding, cells upregulate markers which are associated with intestinal regeneration such as CLU, KRT7, as well as YAP1 and various of its target genes, such as ANXA1 and AREG (Figure 4E)^6,7^. Together with the imaging data (Figure 2), our results indicate that cells pass through an initial phase which mimics damage-induced regeneration of the human intestine, similar to the early phase of mouse intestinal organoid development^6,8,9^. Subsequently, cells start to express the first markers associated with the homeostatic intestine. We identified two major subpopulations of stem cells (SC *Start* and SC *Bal)* which are exclusive to the two applied media and differ in their proliferation state and the expression of markers such as LGR5, ASCL2 (SC *Start*) and OLFM4, KI67 (SC *Bal* (Figures 4C to 4E). Importantly, both SC states are largely negative for YAP1 target genes and thus, do not represent a YAP1^+^ state (Figure 4E).

Secretory progenitors (DLL1^+^) are established from both populations (Sec. 1 and Sec. 2, respectively) and branch out further into the EEC lineage via expression of FOXA2^46^ or towards MUC2^+^/ITLN1^+^ GC by going through progenitors that express ATOH1^+^/SPDEF^+^^47,48^. FABP1^high^ absorptive cells show a separate branch and are exclusive to a trajectory through SC *Bal*, likely as a result of changes in the applied *Balance* medium (Figure 4D). Furthermore, due to the improvements of our medium regime and analysis workflow we were able to observe rare transitions. For example, we observed the transition of secretory progenitor cells (Sec.1 and 2) towards EEC progenitors (EE-pr.) and subsequently observed a bifurcation into two distinct EEC lineages: TPH1^+^/serotonin-secreting EE-non enterochromaffin (EE non-EC) and GCG^+^ enterochromaffin (EC) cells (Figures 1L, 4D and 4E). These are the only two EEC lineages reported in human colonic tissue^49^. Thus, this rich data set will be able to provide a detailed understanding of the developmental trajectory and lineage commitment of human colonic cell types in the context of colonic organoid maturation.

### Organoid regeneration is driven by dedifferentiation trajectories

The unbiased outgrowth from single cells and subsequent organoid development represents a distinct aspect, as it does not involve active selection for specific mature cell types before seeding. The imaging-based pseudotime trajectory shows that organoids express traces of mature cell type markers, such as MUC2/ITLN1 and FABP1 at very early stages before YAP1 upregulation, indicating characteristics of dedifferentiation processes (Figures 2F and 2G, S2F, S5A and S5B).

To further understand this process and the onset of the regenerative response, we subclustered the ‘Regenerative’ cluster in the colonic organoid scRNA-seq to find subpopulations (Figures 5A and 5B). We captured cluster-specific transcriptomic signatures of either colonocytes, exocrine/enteroendocrine cells, or stemness genes and named these clusters ‘Colonocyte reprogramming’ (C rep.), ‘Secretory reprogramming’ (Sec. rep.), and ‘Stem cells reprogramming’ (SC rep.), respectively (Figures 5A and 5B). To infer the similarity of these cells to other populations, we performed diffusion map analysis and trajectory inference, which indicated connections of these reprogramming cells to their mature counterpart and showed that all reprogramming cells go through a YAP1^+^/CLU^+^ state to then re-differentiate into all mature cell types (Figures 5C and 5D). Furthermore, these reprogramming clusters are accompanied by distinct patterns of gene expression. Reprogramming secretory cells upregulate TP53 and the YAP1 target gene CLU, while SC-associated cells exhibit an upregulation of YAP1, TP53, and IL33 consistent with previous observations in mouse models following radiation injury^50,51^. In contrast, reprogramming colonocytes display an upregulation of AREG, PLAUR, and CXADR^52,53^(Figure 5B). Subsequently, cells from all dedifferentiating clusters traverse a distinct cell state represented by the YAP1^+^/CLU^+^ cluster exhibiting minimal expression of genes associated with mature cell types. Instead, genes associated with regeneration markers, such as ANXA1 and DUSP5, as well as YAP1-target genes, including CLU and AREG, are highly expressed within this cluster (Figure 5B).

**Figure 5.**
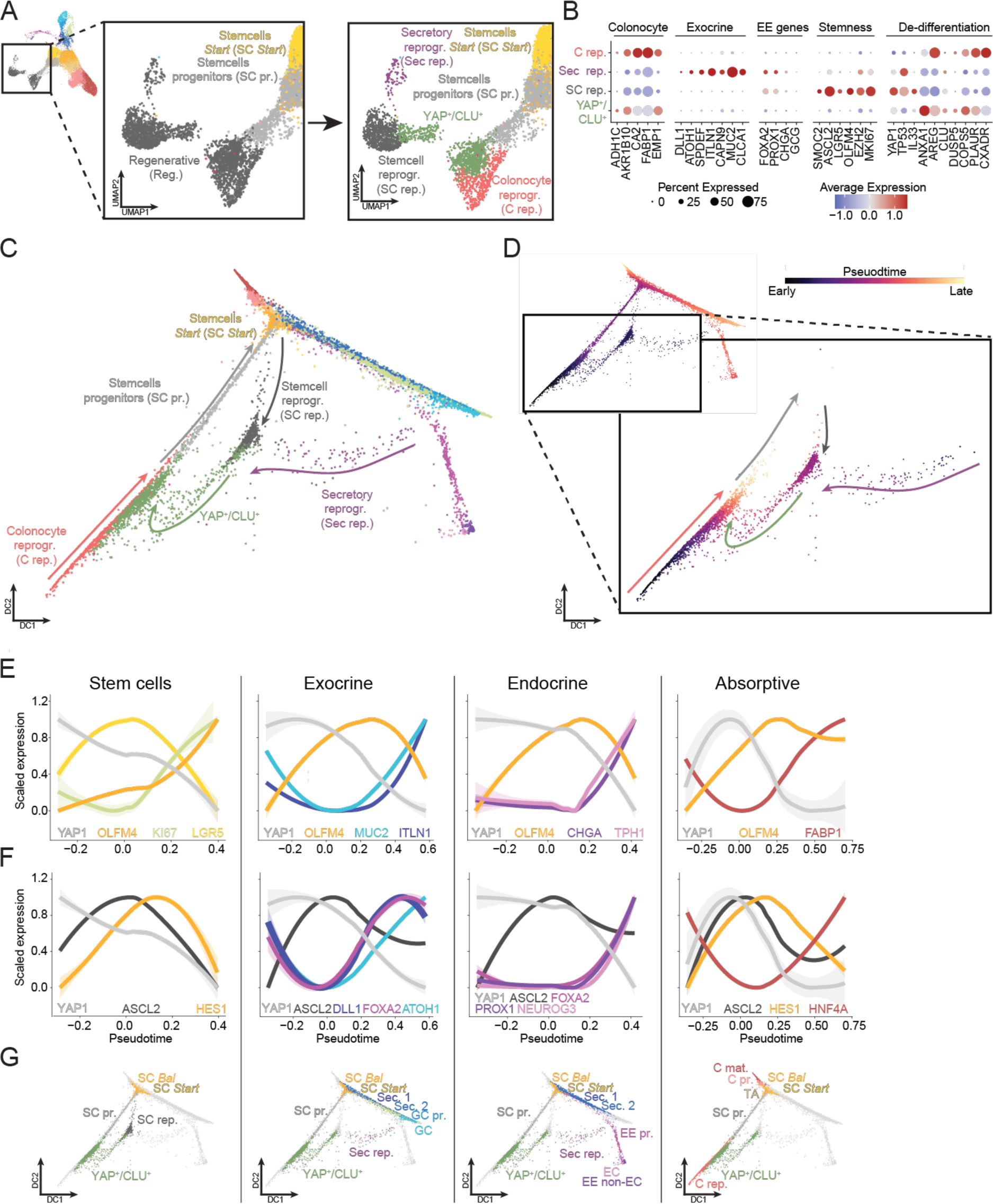
Organoid regeneration is driven by dedifferentiation trajectories. (**A**) Higher resolution annotation of the Regenerative cell-cluster (gray, left) from the time course scRNA-seq data into multiple dedifferentiating subpopulations (right). (**B**) Dot-plot of genes associated with dedifferentiation, stemness, endo/exocrine cells, and colonocytes across refined grouping within the initial Regenerative cluster from the scRNAseq time course. Dot-color shows the average expression level while its size indicates the fraction of positive cells. (**C-D**) Scatter plot of the first two diffusion components of a diffusion component analysis (DCA) of the cell-cycle corrected and sketched scRNA-seq time course data, color-coded by cluster annotations. Arrows indicate directionality within early dedifferentiating cells as indicated by URD pseudotime analysis colored by the representing cluster (**D**, see Methods). (**E-G**) Line plots of expression changes over pseudotime within the stem cell, exocrine, endocrine, and absorptive cell maturation trajectories. Data depicts mean and the 95% confidence interval. Established colonic cell-type marker genes (**E**) and transcription factors (**F**) are shown in relation to YAP1 and OLFM4 (**E**) and YAP1 and ASCL2 (**F**). Used populations for pseudotime computation within indicated lineages are indicated in (**G**).

To study the full maturation trajectory to all mature cell types we performed cell type-specific pseudotime analysis (Figures 5D-5G) and observed a similar transient response of YAP1 expression across all lineages, similar to our protein-level analysis (Figures 2F-2H). To reinforce this observation, we computed a YAP-target gene score, revealing specific upregulation in the YAP^+^/CLU^+^ cluster (Figure S5C and S5D)^7,8^, while the WNT-target gene score exhibited an opposing pattern (Figures S5E and S5F)^54^. OLFM4, on the other hand, showed distinct dynamics across lineages. SC show an increased expression towards the end of the trajectory, whereas secretory cells show an intermediate peak during the stem cell phase of their maturation and subsequent downregulation after secretory specification. Absorptive cells end in a comparatively high steady state expression of OLFM4, suggesting that absorptive progenitors maintain its expression longer, likely through an OLFM4^high^ TA population. Finally, we showcase transcription factor (TF) dynamics of known players in intestinal lineage determination (reviewed by Beumer and Clevers^11^), such as ASCL2^55,56^ and HES1^57^ for SC, DLL1^58^ and FOXA2^46,59^ for exocrine cells, NEUROG3^60,61^ and PROX1^62^ for endocrine cells, and HES1/HNF4A^63,64^ for absorptive cells, and revealed their temporal expression patterns for the first time in a human colonic system along their differentiation trajectory (Figure 5F). Collectively, these findings emphasize the plasticity and dynamic nature of cell states during the regeneration and maturation of organoids.

### Colonic organoids resemble cell types of the *in vivo* tissue

To evaluate the resemblance of our human colonic organoid cultures to *in vivo* tissue, we conducted a comparative analysis with four independent *in vivo* scRNA-seq studies^12,65–67^. Initially, we grouped organoid-derived cells into four major lineages (stem cells (SC), goblet cells (GC), enteroendocrine cells (EEC), and colonocytes (C) (Figure 6A) and compared them to the same granularity of cell type annotation in either normal (’Adult’), inflamed and adjacent uninflamed, or fetal (6-10 (first trimester) and 12-17 (second trimester) post-conception weeks) *in vivo* colonic tissue. This revealed high transcriptional similarity of *in vitro* cell types to their *in vivo* counterpart, especially to normal adult tissue (Figure 6B). We subsequently embedded all the *in vivo* and *in vitro* data into a unified representation (collectively 155,670 single cells), showing notable overlap between datasets from *in vivo* origin (Figure 6C, left panel) and this *in vitro* study (Figure 6C, right panel). After cluster annotation and marker expression profiling (Figure 6D), the only cluster which is not detected in our *in vitro* cultures is a group of cells exclusive to the ‘Fetal’ data set^65^ with elevated CLU expression and void of any canonical cell type markers (Figures 6C and 6D). Additionally, a small population of *in vitro* cells are clustered with the *in vivo* BEST4^+^ cluster.

**Figure 6.**
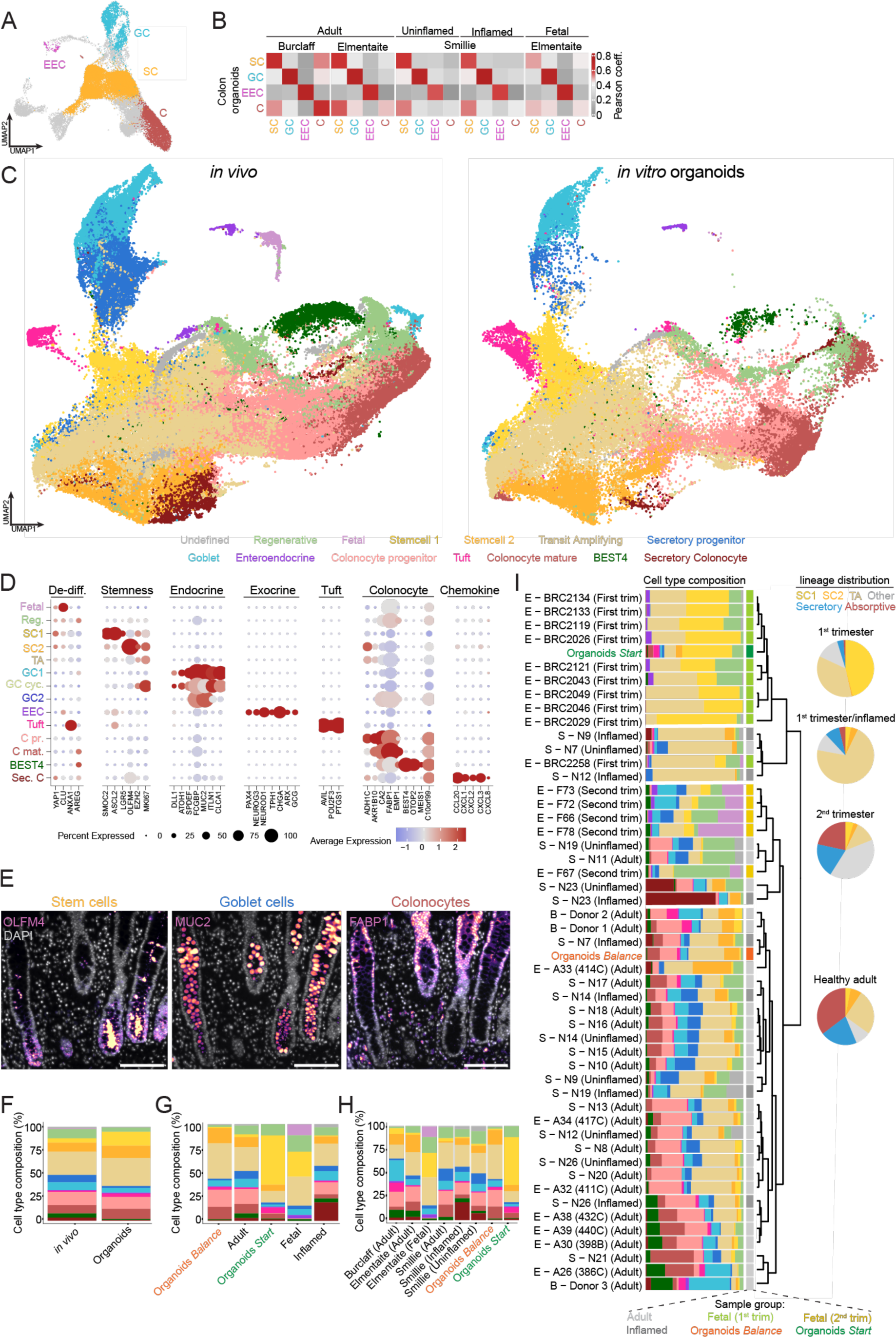
Colonic organoids resemble cell types of the *in vivo* tissue. (**A**) Cell-cycle corrected and sketched time course scRNA-seq data annotated into four broad classes: stem cells (SC), goblet cells (GC), colonocytes (C), and enteroendocrine cells (EEC). (**B**) Heatmap to depict the Pearson coefficient between broadly annotated cell types of the colon organoid *in vitro* data and three similarly binned *in vivo* scRNA-seq data sets (see Methods) of either healthy adult tissue (Adult), uninflamed and inflamed tissue of inflammatory bowel disease (IBD) patients (Uninflamed and Inflamed, respectively), as well as a data from the first two trimester of human fetal development (Fetal). (**C**) Embedding of all *in vivo* and the colonic *in vitro* data from this study into a unified UMAP representation (collectively 155,670 single cells), highlight cells from *in vivo* origin (left) and *in vitro* origin (right). Cells are color coded by a refined cluster-annotation based on the expression profile of individual clusters (see Methods). (**D**) Dot- plot of genes associated with dedifferentiation, stemness, endo/exocrine cells, as well as tuft cells, colonocytes and chemokine expression patterns across annotated clusters from the integrated *in vivo/in vitro* scRNA-seq data. Dot-color shows the average expression level while its size indicates the fraction of positive cells. (**E**) Representative images of primary staining markers used in this study for the stem cell compartment (OLFM4, left), goblet cells (MUC2, middle), and colonocytes (FABP1, right) in colonic tissue slices from biopsies of healthy individuals (see Methods). Images are maximum intensity projections (MIPs) of z-stacks acquired on a widefield microscope. Scale bar, 100 µm. (**F-I**) Cell-type composition of scRNA-seq samples of binned either by organoids (this study) and all merged *in vivo* tissues (**F**), their *in vivo* categories and the two organoid media conditions (**G**), per individual study (**H**) and per individual (**I**). Pie charts in (I) presents a simplified average lineage distribution among major branches found by the dendrogram. Annotations are based on the merged data set of (C). E: Elmentaite et a l., (2020 and 2021), S: Smilie et al. 2019, B: Burclaff et al., 2022, trim : trimester.

Furthermore, to probe not only the molecular similarity but also the spatial organization of the epithelial cells, we used colon tissue slices of adult individuals to identify the *in vivo* localization of major protein markers used in this study (Figures 6E and S6A). Similar to the localization observed *in vitro* (Figures 2H and S2G), OLFM4 is secreted and preferably localizes towards the crypt-bottom while FABP1^+^ cells are localized towards the upper-half of the crypt. MUC2^+^ GCs and both subtypes of EECs are interspersed throughout the full epithelium.

Additionally, we investigated the cell type composition of organoids and *in vivo* tissues (merging all datasets, Figure 6F), in different *in vivo* categories and the two organoid media conditions (Figure 6G), per individual study (Figure 6H) and per patient (Figure 6I). We observed considerable interpatient differences in cell type composition from *in vivo* sequenced tissue. Our *Balance* medium generates cell types in a proportion similar to the *in vivo* tissue of normal adult individuals (Figures 6F to 6H). Observed differences may emerge from the exact origin of the sequenced tissue along the colonic proximal-distal axis^14^ or technical variance. However, the organoid cell type composition falls within the patient-to-patient variability.

In contrast, organoids cultured in the *Start* medium display a cell type composition reminiscent of first-trimester colonic samples, distinct from adult individuals due a substantial SC1 population (Figures 6G to 6I). SC1 is the dominant SC state observed in fetal samples, indicating that SCs of human colonic organoids transiently have a transcriptomic signature similar to SCs found in fetal colonic development. Thereafter, a more mature cellular composition is established in the *Balance* medium containing mostly SCs akin to adult individuals represented by the SC2 population (Figure S6B). When examining the clusters derived from the integrated representation (Figure 6) alongside those from the isolated *in vitro* UMAP (Figure 4), it becomes evident that *in vitro* cells in the SC1 population primarily belong to the *in vitro* SC prog. and SC *Start* clusters, whereas cells in SC2 are found in the *in vitro* SC *Bal* and TA clusters (Figure S6C).

This analysis emphasizes the resemblance of mature organoids to adult tissue in terms of transcriptional identity, cell type proportions, and their spatial distribution. Additionally, we report the dynamic nature of the SC compartment as regenerating organoids progress from a fetal-like to an adult-like SC state.

### An initial fetal-like stem cell state enhances intestinal regeneration

The developed workflow presents an opportunity to improve our understanding on the dynamics of the SC states and their functional implications in a human model system. These SC states are represented by two distinct clusters (SC1 and SC2) in the integrated data set (Figures 6C and 6D) as well as primarily two clusters (SC *Start* and *SC Bal*) in the stand-alone *in vitro* data (Figures 4D and 4E).

We first tracked the expression of SC marker genes over pseudotime in our scRNA-seq data, separated by the applied medium (Figure 7A). In the *Start* medium, ASCL2 and LRG5 are upregulated, followed by their downregulation in the *Balance* medium. OLFM4 shows a similar initial increase in expression in the *Start* medium but is transiently downregulated with a second peak in expression towards the end of the *Balance*-medium SC trajectory (Figures 4F and 7A). The switch from an ASCL2^+^/LGR5^+^ (SC *Start*) to an OLFM4^high^ (SC *Bal*) profile is accompanied by a switch from NOTCH1 to NOTCH2 expression, in line with a study on SC maturation trajectories from *in vivo* fetal to adult tissue in the human intestine (Figure 7B)^68^. This further underscores the fetal-like transcriptomic signature observed in SC *Start* before an adult-like SC *Bal* state is established.

**Figure 7.**
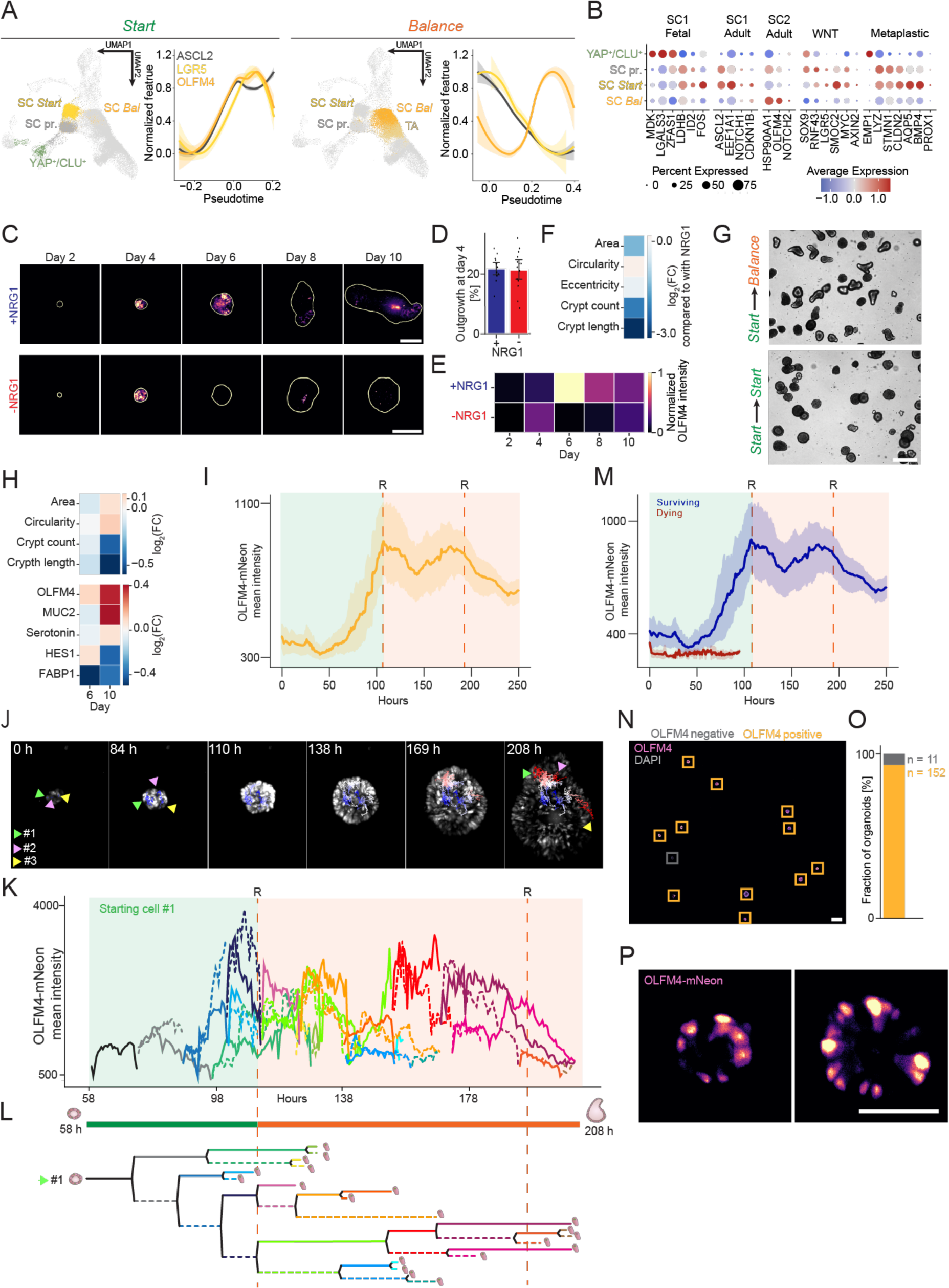
An initial first trimester-like stem cell state enhances intestinal regeneration. (**A**) URD pseudotime analysis (see Methods) of the stem cell trajectory divided by samples taken within the first four days of organoid maturation during *Start* medium incubation (left) and during the last six days of organoid maturation during *Balance* medium incubation (right). Data shows 95% confidence interval. (**B**) Dot-plot of genes dominantly expressed in *in vivo* fetal or adult cells of the SC1 cluster as well as of cells in the *in vivo* adult SC2 cluster compared two four regenerative and SC states of the *in vitro* data. Genes were defined by pairwise differential expression analysis. Moreover, genes associated with WNT-pathways activation and a metaplastic signature are shown. Dot-color shows the average expression level while its size indicates the fraction of positive cells. (**C**) Representative images of OLFM4 marker expression in organoids along fixation timepoints in control medium with NRG1 (top), as well as medium devoid of NRG1 (bottom). White outline represents the segmentation mask. Images are maximum intensity projections (MIPs) of z-stacks acquired on a spinning-disk confocal system. Scale bars, 100 µm. (**D**) Outgrowth quantified at day 4 with of organoids grown with (left) or without (right) NRG1. Data denotes mean and the 95% confidence interval. (**E**) Log2(fold change) of morphological features of mature organoids grown with NRG1 supplementation compared to without. (**F**) Heatmap of normalized OLFM4 mean intensity along five sampling timepoints grown with (top) and without (bottom) NRG1. (**G-H**) Comparison of organoids grown in control conditions (*Start* to *Balance* medium switch at day 4), or exclusively in Start medium. (**G**) Representative brightfield images of organoids in the control condition (top), and organoids grown purely in *Start* (bottom) medium. Scale bars, 100 µm. (**H**) Heatmap depicting log2(fold change) of morphology (top) and mean intensity of markers for cell types (bottom) of organoids grown purely in *Start* medium compared to the control. Two timepoints (D6 and D10) are shown. Related to Figure S7A. (**I**) Quantification of organoid-level OLFM4-mNeon intensity along the organoid maturation trajectory on a light sheet microscope over 250 hours. Green background indicates timeframe in *Start* medium, orange background time in *Balance* medium. R and dashed lines indicate timepoints of medium refreshments. Data depicts mean and 95% confidence interval. (**J-L**) Quantification of cell-level OLFM4-mNeon intensity along the organoid maturation trajectory on a light sheet microscope between 58 and 208 hours after seeding of single cells (**K**). Shown is the lineage of starting cell 1 out of 3 (related to Figures S7F and S7G). Green background indicates timeframe in *Start* medium, orange background time in *Balance* medium. R and dashed lines indicate timepoints of medium refreshments. (**L**) Cell-lineage tree for highlighted cells in (**J**, starting cell 1 out of 3) including snapshots of the tracked organoid on a light sheet microscope at the indicated timepoints with positions of all three traced starting cells (cell 1: green, cell 2: pink, cell 3: yellow). Tracks are colored and stylized based on the cell id. Tracks which were lost are cells with expression levels of OLFM4 below the detection threshold, or cells with unclear movement of the cell. (**M**) Quantification of organoid-level OLFM4-mNeon intensity along the organoid maturation trajectory color-coded by the fate of the organoid (surviving: blue, dying: red) on a light sheet microscope over 250 hours. Green background indicates timeframe in *Start* medium, orange background time in *Balance* medium. R and dashed lines indicate timepoints of medium refreshments. Data depicts mean and 95% confidence interval. (**N-O**) Quantification of OLFM4^+^ positive organoid fraction after initial outgrowth. Representative image of a full-well of the OLFM4-P2A-mNEON-NLS reporter line at day 4. OLFM4^+^ organoids are indicated with orange rectangles, OLFM4^-^ with grey rectangles. Images are maximum intensity projections (MIPs) of z-stacks acquired on a spinning-disk confocal system. Scale bar, 50 µm. (**O**) Quantification of (N). (**M**) Representative images of two organoids of the OLFM4-P2A-mNEON-NLS reporter line at day 4. Images were acquired on a light sheet system. Scale bar, 50 µm.

Next, we aimed to understand how both observed SC states are induced in the *Start* and *Balance* medium, respectively. This medium switch is accompanied by a decrease in EGF and WNT-signaling, as well as increased NRG1-signaling (Table 3). As ASCL2 and LGR5 are WNT-target genes and highly expressed in the *Start* medium, we probed a general WNT- signature (SOX9, RNF43, SMOC2, MYC, AXIN2) across pseudotime in the SC trajectory. We observed high activity of the WNT-pathway in the SC *Start* population until the switch to the *Balance* medium, indicating that elevated WNT-activation correlates with the SC *Start* state (Figure 7B, S5E and S5F). We hypothesized thereafter that the switch to the SC *Bal* population is associated with increased NRG1-signaling. We assessed this hypothesis in an imaging time course over 10 days without NRG1 supplementation (Figures 7C-7F). To read out the current SC state in our organoid cultures, we made use of the unique expression profile of OLFM4, a stem cell marker with overlap in both SC states. As mentioned before, it exhibits an initial peak around day 3-4 upon entering the SC *Start* state and a second peak around day 6 when the SC *Bal* state is established and first crypts form (Figures 4F, 7A). We observed that organoids without NRG1 survive in unchanged efficiency (Figure 7D) and induce the initial OLFM4 expression linked to entry into the SC *Start* state (Figures 7C and 7E). However, the entry into the SC *Bal* state is interrupted as no second OLFM4 peak is observed (Figures 7C and 7E) leading to failure of undergoing crypt-morphogenesis and correct spatial organization of cell types (Figure 7F).

Our focus shifted to understand the functionality of both SC populations. Given the strong correlation between the applied medium (i.e., *Start* or *Balance*) and the occurrence of the two SC-states (Figures 4C and 4D), we aimed to enrich for SC *Start* and SC *Bal* states by culturing organoids exclusively in *Start* and *Balance* media, respectively (Figures 7G, 7H and S7A). Organoids grown solely in *Balance* medium (supplemented with the essential ROCK-inhibitor for initial outgrowth) exhibited reduced viability and atypical phenotypes (Figure S7A). In contrast, utilizing only *Start* medium resulted in expected formation efficiency from single cells, but organoids characterized by decreased crypt-morphogenesis and bias towards secretory and OLFM4^+^ cells as observed in high-content microscopy (Figures 7G and H) and validated by FACS (Figure S7B). This suggests that an initial population of SC *Start* is linked to an efficient regenerative process, while a switch towards SC *Bal* allows efficient differentiation into the absorptive lineage and continuous support of crypt-morphogenesis.

To delve into these aspects with more temporal resolution, we engineered an OLFM4-P2A- mNeon-2xNLS reporter line (Figures S7C and S7D, Methods)^65,66^. Light sheet live-imaging of 11 organoids from single cells until maturation at 250 h with 30 min sampling intervals revealed detailed organoid-level dynamics of OLFM4 (Figures 7I and S7E, Video S3). In the *Start* medium, OLFM4 expression increases steadily in almost all cells in the organoid indicating entry into the SC *Start* state for all cells at that time. Subsequently, when the *Balance* medium is applied, there is a transient downregulation of OLFM4, associated with the shift towards the SC *Bal* population, aligning with our gene expression data (Figures 4F, 7A). Tracking of 39 cells starting from three OLFM4^+^ cells during organoid growth between 58 and 208 h shows cells with variable OLFM4 intensity profiles (Figures 7J-7L, S7F and S7G). While the almost all cells show an initial upregulation of OLFM4 before the medium switch, only a subset of tracked cells enter the second peak in expression associated with the SC *Bal* state (Figures 7J-7L, S7F and S7G, Video S4), suggesting a requirement to enter the SC *Start* state to conclude regeneration, but the high plasticity of this state allows to differentiate into multiple lineages thereafter. Similar to our observations during light sheet imaging of the H2B-iRFP670 line (Videos S1 and S2), we again found that 3 (27.3%) out of 11 organoids did not survive the long-term imaging (Video S5). Evaluation of OLFM4 dynamics in these organoids revealed that all organoids, which did not induce initial OLFM4 expression and, thus, the emergence of SC *Start* cells during the initial regenerative response, died around the same time (91, 98 and 101 h, Figures 7M, S7H and S7I). To validate this, we quantified the relative abundance of OLFM4^+^ organoids on day 4 after the regenerative phase is largely concluded (Figures 7N and 7O). At this stage, almost all surviving organoids (∼93.3%) displayed OLFM4 expression (Figure 7O) with a significant proportion of cells within an organoid being positive for this marker (Figure 7P), resembling the SC-biased cell type composition in the scRNA-seq of colonic tissues during the first trimester (Figure 6I). As the observed initial increase in expression until day 4 is linked to the establishment of the SC *Start* state (Figures 4F and 7A), it once more highlights that entry into this population concludes a successful regeneration.

In summary, the fetal-like SC *Start* population plays a pivotal role in the successful regeneration and cellular plasticity of colonic organoids and is in the context of our workflow likely induced through elevated WNT-activation in the *Start* medium. Successful establishment of subsequent the adult-like SC *Balance* state is associated with NRG1 supplementation and reduced WNT-signaling, leading to continuous support of crypt-morphogenesis.

## Discussion

The regeneration of the human colon is a complex and fast-paced process. *In vivo* studies typically provide only a snapshot of the regenerative state, and until now, no study has investigated the immediate response of adult human intestinal cells after damage comprehensively. Our developed workflow fills this crucial gap by providing a highly dynamic platform to study both cell- and tissue-level responses to damage. Resulting organoids of this workflow can be kept long-term in culture and resemble the *in vivo* tissue remarkably well in terms of cell type composition, the spatial patterning of cell types, and their transcriptomic signature.

Our workflow allows outgrowth from a single cell and thus provides a streamlined and robust workflow to probe therapeutical avenues in a human setting. We utilized this with therapeutically utilized inhibitors against ERBB-receptors and activating EGF-like ligands to enhance our understanding of the complex ERBB-receptor signaling network in the intestinal epithelium. We report that initial survival after damage, as well as crypt-morphogenesis in the transition from regeneration toward homeostasis, is connected to active EGFR-homodimer signaling. However, our results could not explain the low efficacy of ERBB3 inhibition compared to inhibition of its heterodimerization partner ERBB2, or removal of its ligand NRG1. While we and others did not observe expression of ERBB4, the second known receptor for NRG1, its expression could be context dependent. For example, it is shown to be overexpressed in pathological contexts such as necrotizing enterocolitis (NEC)^70^, IBD^71^, and CRC^72–74^ while no expression was detected in healthy adult, homeostatic tissues^37^. Given that the inhibition of ERBB2 had the most pronounced effects during the regenerative phase, we hypothesize that ERBB4 may be expressed as a backup-mechanism transiently after damage during this period. Furthermore, reports of the ERBB2 inhibitor Pertuzumab indicate varying efficacy in blocking its heterodimer binding partners compared to inhibition of homodimers^39^. This phenomenon may extend to the ERBB3 inhibitor Lumretuzumab.

We then benchmarked the overlap of the developed *in vitro* system to publicly available scRNA-seq data sets of healthy fetal, adult, or inflamed *in vivo* tissues. To our surprise, we noted only limited overlap between the transcriptomic signature of individuals with IBD and the transient regenerative state of our *in vitro* model, likely due to the chronic nature of the disease in combination with the missing immune system in our model^69^. However, our study identified a distinct stem cell population (SC *Start*) which arises during the fourth day after single cell seeding shortly before first mature cell types are observed. These cells exhibit transcriptomic characteristics of SC occurring during fetal colonic development and notably lack the expression of YAP1 target genes. Therefore, we hypothesize that these cells are likely to conclude the regenerative process. The transition to a fetal-like state following damage has been reported in mice^4,6,50,75^ and recently in a human *in vitro* and *in vivo* study as a crucial step in metastatic growth during colorectal cancer (CRC) progression^76,77^. Indeed, comparison of our *in vitro* study to an *in vivo* dataset of metastatic CRC progression revealed similarities between the SC *Start* state and stem cells of primary and metastatic tumor tissue (data not shown)^76^. Within our system, SC *Start* represents a brief bottleneck, in contrast to the prolonged presence of related fetal-like stem cell states in both cancerous and developmental contexts. We argue that during development and disease progression, this state is stabilized through signals of the local microenvironment, mechanical feedback, and mutational burden in the case of cancer. *In vitro*, a switch to medium with lower EGFR and WNT signaling leads to sudden loss of this population. Likely not coincidently, mutations of the EGFR (KRAS^G12D^) and WNT (APC^-/-^) pathways are the two most common oncogenic drivers in CRC^78^. The gradual downregulation of fetal and metaplastic (Figure 7B) features towards maturing SC *Start* and SC *Balance* populations emphasizes the significance of cellular plasticity during the regenerative process. This intriguing observation, in combination with the accessibility of the *SC Start* state and scalability of our model system, may open new possibilities for druggable targets in the context of intestinal regeneration and CRC^79,80^. However, it is crucial to recognize that this strategy embodies a dual nature. Targeting the fetal-like population in CRC disease progression could provide important therapeutic avenues, but it may hinder tissue regeneration after damage from radiation and chemotherapy by obstructing the transition through the highly similar and vital *SC Start* state.

Although robust, the seeding of single cells as an experimental start timepoint in our workflow may represent a severe method of damage modeling. To further expand our understanding, future investigations should explore how the human intestine responds to other sources of damage, such as SC ablation, mechanical disruption of mature organoids, or exposure to chemical toxicity. Additionally, patient-derived organoids of individuals of different ages, colonic location, and those with diseased tissue, such as CRC and IBD, may provide further insights into the regenerative process and its deregulation in pathologies.

In conclusion, our study provides a detailed examination of how the human colon responds to extensive damage in an *in vitro* model, offering insights at the protein and mRNA levels at high temporal resolution. Our results highlight the remarkable plasticity of intestinal cells, likely as an adaptive response to the harsh environment in which they thrive. These findings advance our understanding of the regenerative processes in the human colon and offer potential avenues for therapeutic intervention in intestinal disorders and diseases.

## Limitations of the study

While our study has yielded valuable insights into human colon organoid regeneration and stem cell regulation, it is important to acknowledge its limitations. First, our research primarily relies on an *in vitro* organoid model, which, while highly informative, may not fully recapitulate the complexity of the *in vivo* colonic environment. Additionally, our study focuses on normal organoid development, and the applicability of our findings to disease contexts requires further investigation. Another limitation is the use of a culture medium, which may influence cell phenotypes and introduce potential artifacts. Furthermore, our study predominantly explores the dynamics of organoid maturation within a defined time frame, and longer-term investigations may reveal additional insights into the stability of cell states and phenotypes. Finally, while we have identified potential therapeutic targets and implications, further experimental validation is necessary to translate these findings into clinical applications.

## Acknowledgments

We thank E. Tagliavini for IT support; F. Moos for assistance for light sheet microscopy data; S. Aluri for sequencing; H. Köhler for FACS assistance; D. Gaidatzis for input on scRNA-seq analysis; I. Lukonin, S. Reither and L. Gelman for discussions regarding imaging and screening; E. Tagliavini and N. Repina for discussions on image analysis; the Foundation Hubrecht Organoid Technology (now Foundation Hubrecht Organoid Biobank, https://www.hubrechtorganoidbiobank.org/) for providing all organoid lines used in this study, especially the main colon line used in this study (HUB-02-A2-040); We thank colleagues and laboratory members for scientific discussions and critical feedback on the manuscript. Funding: This project has received funding from the European Union’s Horizon 2020 research and innovation programme under grant agreement No 874769 and Chan Zuckerberg Foundation Grant CZF 2019-002440. This publication is part of the Human Cell Atlas: https://www.humancellatlas.org/publications.

## Author contributions

Conceptualization, K.C.O., M.K. and P.L.; Methodology, K.C.O. and M.K.; Investigation, K.C.O., M.K., G. d. M. and S.Su.; Formal Analysis, M.K. and S.B.; Writing, K.C.O., M.K. and P.L.; Visualization, K.C.O., M.K. and S.B.; Resources, L.C.M. and V.K., Supervision, P.L., S.Sm. and M.B.S.; Funding Acquisition, P.L.

## Data and code availability

The data in this study is available upon request from the lead author if the legal framework permits. Code used for image analysis in this work is publicly available at GitHub: https://github.com/fmi-basel/HT_OrganoidAnalysis

## Methods

### Patient-derived human organoid lines

Tissue material was originally obtained from patients included in HUB-Cancer protocol (12- 093). Patient samples are collected with freely given, specific, informed and unambiguous consent and ethics approval for organoid derivation and data collection (Biobanks Review Committee, UMC Utrecht, sub-bank numbers 12/093). Used organoid lines (see table below) in this human study were provided by the Foundation Hubrecht Organoid Technology (now Foundation Hubrecht Organoid Biobank) (https://www.hubrechtorganoidbiobank.org/). Most experiments were conducted on HUB-02-A2-040 (human colon) or HUB-04-A2-001 (human small intestine).

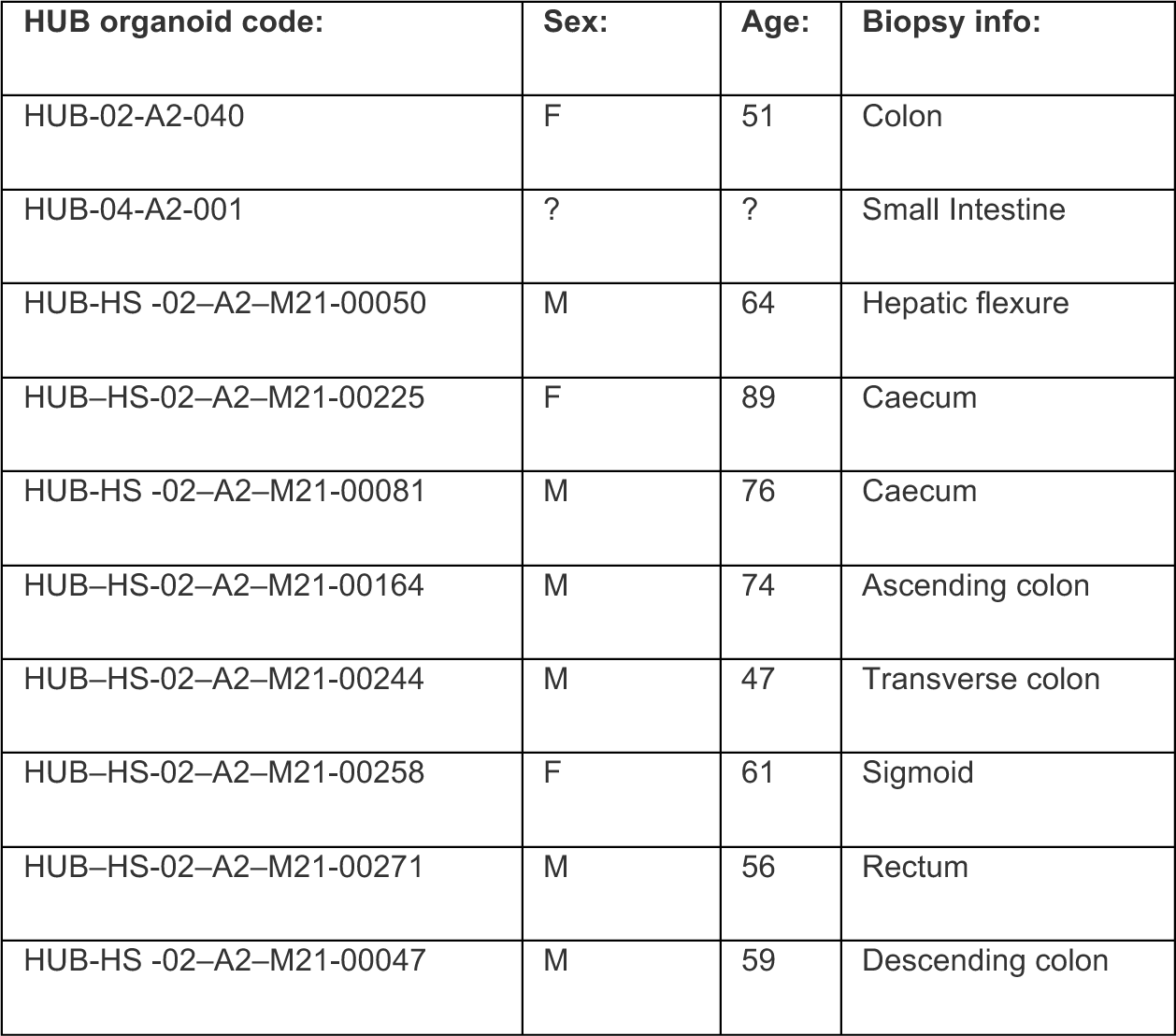

### Human intestinal organoid maintenance cultures

Mature organoids are collected by rinsing the culture well with 1-2 ml ice cold DMEM++ (= DMEM/F12 + 15 mM HEPES (Stem Cell Technologies) + 1x GlutaMAX (Thermo Fisher Scientific) + 100 U/ml Penicilin/Streptomycin (Thermo Fisher Scientific)) and transferring the solution into a 15 ml tube (Falcon). Next, DMEM+++ is added until a total volume of 7 ml is reached before centrifuging the tube at 400 rcf for 4 min at 4°C. Supernatant is discarded and organoids are resuspended into 2 ml of prewarmed 0.05% Trypsin + EDTA (gibco) and incubated for ∼10 min at 37°C. Trypsinization is stopped using 2 ml of a trypsin-inhibitor (Thermo Fisher Scientific) as well as 3 ml of DMEM+++ after reaching a single-cell suspension.

Singularized organoids are then centrifuged at 600 rcf for 5 min at 4°C and supernatant is discarded subsequently. Next, the cell pellet is resuspended into *Start* (Table 3) medium and then filtered through a 40 μm filter to remove large debris and cell clumps. Filtered cells were FAC-sorted to remove remaining smaller debris as well as doublets and count the number of remaining cells.

Finally, the cell-suspension is diluted to 20-35 cells/μl into a ∼66% Matrigel (corning): ∼34% *Start* (Table 3) mixture. Here, lower seeding densities between 20 and 35 cells per μl are recommended as very dense plating affects the morphology and growth of organoids negatively. The optimal seeding density needs to be optimized for every line. For maintenance, the Matrigel mixture is plated into cell culture plates (Corning) in 15-20 μl droplets and are covered after a 20 min 37°C solidification period with warm *Start* medium.

Medium is changed after 4 days to *Balance* (Table 3) and is refreshed once with new *Balance* medium at day 7 until maturation is reached, typically at day 10 for colonic organoids. Small intestinal organoids showed slower growth rates and reached maturation at ∼day 13. If mechanical splitting of mature organoids is desired, it is possible, but one must maintain the culture in *Start* medium for 1 day post-splitting, followed by subsequent medium changes every other day using *Balance* medium.

### Sample seeding into imaging plates

#### 96-well imaging plates

Trypsinized cells were first resuspended into a ∼66% Matrigel (corning): ∼34% medium (see experimental methods for further details) mixture with a density of 35 cells/µl unless stated otherwise. A 5 µl drop was placed centrally into a pre-warmed 96- well imaging plate (Greiner #655090) and overlayed with 70 µl medium after a solidification period of 20 min at 37°C and placed into a 5% CO2, 37 °C incubator thereafter. Due to edge-effects, all border wells were excluded from experimental workflows and instead filled with PBS.

#### 384-well imaging plates

384-well plates (PerkinElmer #6007550) were first pre-cooled to 4°C and wells were covered with 10 µl of an ice-cold ∼66% Matrigel (corning): ∼34% medium (see experimental methods for further details) mixture containing no cells using an INTEGRA ASSIST PLUS pipetting robot. The mixture was spun down carefully into the well bottom before a slight solidification period of 10 min at RT. In the meantime, trypsinized cells were resuspended into medium (see experimental methods for further details) with a density of 35 cells/µl unless stated otherwise. 40 µl of the cell-suspension was pipetted into all wells with the help of a washer-dispenser robot (BioTek EL406). Single cells were subsequently spun into the Matrigel mixture carefully in such a way that they end up centrally in the Matrigel-layer. Finally, the plate is placed in a 5% CO2, 37 °C incubator thereafter. Due to edge-effects, all border wells were excluded from experimental workflows and instead filled with PBS.

### Sample preparation for imaging

#### Fixation

Samples in imaging plates were first spun for 10 min at 10 °C and 1000 RCF in a plate centrifuge. Subsequently, wells were overlaid with equal volume of 8% PFA in PBS (final concentration: 4% PFA in PBS) and incubated at RT for 30 min. Subsequently, wells were rinsed thrice with PBS and washed twice for 10 min at RT on a shaker with PBS. For experiments which used 384-well imaging plates, samples were usually washed on a BioTek EL406 liquid dispenser.

#### Staining for immunofluorescence

Samples in imaging plates were incubated in a 0.5% Triton X-100, 100 mM NH4Cl, 3% donkey serum in PBS solution for 3h at RT on a shaker to permeabilize the sample and reduce unspecific antibody bindings. Subsequently, samples were rinsed thrice with PBS before incubating them in a primary antibody solution ON at RT. Primary antibodies were diluted in a 3% donkey serum, 0.1% Triton X-100 in PBS solution as follows: anti-OLFM4 1:200; anti-MUC2 1:200; anti-FABP1 1:200; anti-KI67 1:500; anti-CK20 1:200; anti-Serotonin 1:1000; anti-CHRA 1:500; anti-ITLN1 1:200; anti-POU2F3 1:200; anti-NUSAP1 1:200; anti-LYZ1 1:1000; anti-NOTCH1 1:200; anti-LRIG1 1:200; anti-Fabp2 1:200; anti-Agr2 1:500; anti-YAP1 1:200. Unbound and weakly bound antibodies were washed off by rinsing the sample thrice with PBS and washing three times for 10 minutes at RT on a shaker. Samples were then overlayed with secondary antibodies against the used primary species for 3 h at RT on a shaker. For this, secondary antibodies were diluted in the same buffer as primary antibodies at a concentration of 1:500. Additionally, the solution was supplemented with 0.2 µg/ml 4’,6-Diamidino-2-Phenylindole, Dihydrochloride (DAPI, Invitrogen #D1306) to stain DNA. For experiments utilizing 384-well imaging plates, samples were usually washed on a BioTek EL406 liquid dispenser.

### High-content imaging

1 h before start of the imaging procedure, samples were placed into a refractive-index-matching imaging buffer^81^. Subsequently, all samples were imaged on Yokogawa CV7000S automated spinning-disk high-content microscope. To reduce the time and amount of generated data, samples were first scanned with a 405 nm excitation laser and a 4x air objective (Olympus, NA = 0.16) to acquire low-magnification maximum intensity projection (MIP) overviews of wells. MIPs were then used for a threshold-based segmentation workflow with a custom-written ImageJ macro^5^ to identify organoids based on the brightness of the DNA (DAPI) signal with a minimum-size filter, typically set smaller than the size of single cell to remove debris. Finally, the segmentation map was upscaled by a factor of 5 and coordinates of identified objects were used as an input for the subsequent higher magnification imaging with a 20x air objective (Olympus, NA = 0.75). Samples were imaged from 10 µm below the position found by the autofocus until the maximum height of the sample depending on the current organoid state in 5 µm steps. For very large organoids, the imaging was stopped 120 µm in z due to loss of faithful signal. Unless stated otherwise, only 16bit MIPs were saved and used for subsequent analysis workflows.

### EGF-like ligand titration

Before establishment of the final media (Table 3), organoids were grown in an initial medium, influenced by previous reports (Table 4)(^19,26^) and supplemented with 5 ng/ml EGF. However, 4 h before collection, all EGF-like ligands were removed from the medium to reduce bias due to the maintenance medium. Organoids were collected and trypsinized to single cells as described before. Next, cells were resuspended and seeded as mentioned before in either 96- well (NRG1, EREG, AREG supplementation experiments,) or 384-well (time resolved EGF/NRG1 titration, PerkinElmer) imaging plates. The initial medium was supplemented with either 0/50 ng/ml EGF in combination with either 0, 10 or 100 ng/ml NRG1, EREG, or AREG for EGF-like ligand supplementation experiments. For the time resolved EGF/NRG1 titration, cells were resuspended at a density of 20 cells/µl (colonic organoids) or 30 cells/µl (small intestinal organoids) and the medium (Table 4) was supplemented according to Table 1 (colonic organoids) or Table 2 (small intestinal organoids). The medium was refreshed on day 4 and day 7 of growth. On day 10, the sample was fixed and stained with DAPI and secondary immunofluorescence as described above.

**Table 4:**
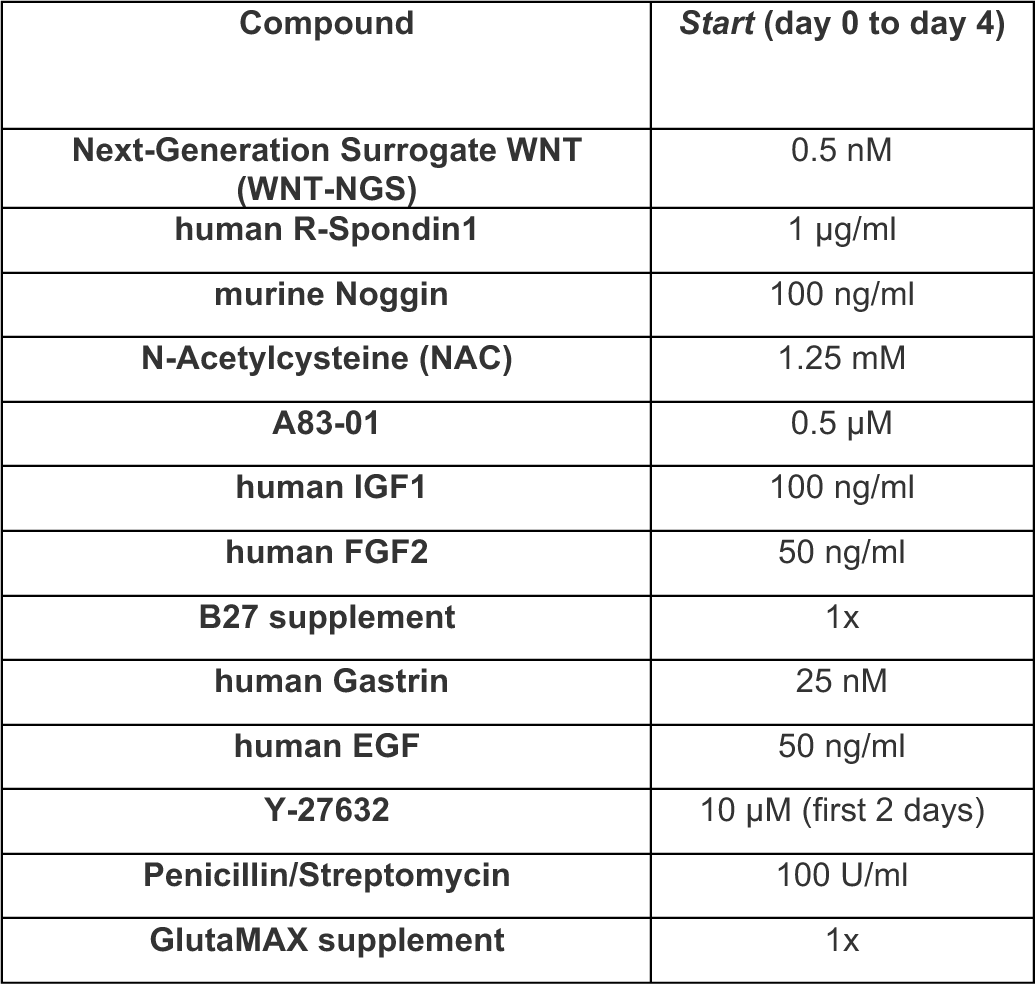
Initial human intestinal medium composition. Composition of the initial medium for the colonic and small intestinal organoids. Ingredients should be diluted with DMEM/F12 (+15 mM HEPES).

### Comparison to other published protocols

Human colonic organoids were grown in the *Start*/*Balance* (Table 3) media and seeded into 96-well imaging plates as described before. However, cells were resuspended into either Intesticult OGM (Stem Cell Technologies), the medium described by Fujii and colleagues^19^, or by a modified version of a medium described by Sato and colleagues^18^. On day 4 of growth control organoids were switched to *Balance* medium, while other conditions were refreshed.

Another refreshment was done at day 7 before the sample was fixed and stained with DAPI and antibodies against and secondary immunofluorescence as described above on day 10.

### Mouse and rat organoid lines and culture

All animal experiments were approved by the Basel Cantonal Veterinary Authorities and conducted in accordance with the Guide for Care and Use of Laboratory Animals. For the experiments, 12-week-old male outbred mice and 24-week-old male Sprague Dawley rats (CD®) were utilized. In the experiments, the following mouse and rat lines were employed: Lgr5::DTR-eGFP (provided by Genentech, de Sauvage laboratory) for mice and CD® (Sprague Dawley) IGS rats, Crl:CD (SD) (kindly donated by NIBR, Novartis) for rats. The small intestine and colon of rats and the colon of mice were longitudinally opened and thoroughly washed with cold phosphate-buffered saline (PBS).

Organoids were generated from isolated crypts of the murine and rat colon as previously described^18^. In summary, segments of the intestinal tissue (including murine and rat colon, as well as rat small intestine) were initially rinsed with cold PBS. Subsequently, the tissue fragments were subjected to a 40-minute incubation in 5 mmol/L EDTA in PBS on a rotating platform at 4 degrees Celsius. After the removal of the EDTA buffer, the tissue fragments were vigorously shaken in 20 mL of cold PBS for 1 minute. This process was repeated two additional times using separate 20 mL PBS-filled tubes. The crypt fractions were examined for morphology, and the cleanest and most suitable fraction was selected for further processing. Isolated crypts were collected, washed with 10 mL of cold DMEM/F12 medium containing 15 mM HEPES, 1x GlutaMAX, and 100 U/mL Penicillin/Streptomycin, and transferred to a 15 mL Falcon tube. After 1 day, the medium was switched to Balance medium (see Table 3) and replaced every 2 days. At this point, the decision could be made between two methods: mechanical splitting or single-cell isolation using 0.05% Trypsin EDTA (Gibco). For mechanical splitting, the culture was maintained in Start medium for 1 day, with subsequent medium changes every other day using Balance medium.

Alternatively, for single-cell isolation, the same preparation protocol as described for human colonic cultures was followed until the final step, where the cell suspension was diluted to a concentration of 150-200 cells/μl in a mixture of 70% Matrigel (Corning) and 30% Start medium (Table 3). After single-cell outgrowth, the medium was changed to Balance medium on day 3 and then replaced with fresh Balance medium on day 5 and day 7, continuing until maturation, which typically occurred between day 7 and day 9 for colonic organoids.

### High-content imaging time course

Human colonic organoids were grown in the *Start*/*Balance* (Table 3) media and seeded into multiple 96-well imaging plates as described before at a density of 35 cells per µl of the Matrigel/*Start*-medium mixture (175 cells/well). A subset of plates was fixed as described above at 48, 96, 144, 192, and 240 h after seeding of single cells. Finally, all plates were simultaneously immunostained as described above and finally imaged on a Yokogawa CV7000S spinning-disk confocal microscope in accordance to aforementioned protocol.

### ERBB-receptor inhibition screen

To test the effects of ERBB-inhibition during different treatment-windows, organoids were initially grown in maintenance medium and seeded into 384-well plates as described above. However, instead of resuspending cells into *Start* medium, cells were resuspended into 20 µl 2x *Start* medium before being seeded at a density of 225 cells/well. The following concentration were tested: Cetuximab 3.5/0.35/0.035 nM, Pertuzumab 25/2.5/0.25 µg/ml, and Lumretuzumab 10/1/0.1 µg/ml. Four types of controls were used with varying amounts of added PBS into DMEM+++ depending on the volume the drugs were dissolved in, in addition to a control without added PBS. Subsequently, 20 µl of the drugs or controls were placed into wells belonging to plates which are used for inhibitor treatment during the initial days (day 0 to 4, and day 0 to 10 treatment windows) in a pseudo-randomized manner to exclude plate- positioning effects using an INTEGRA ASSIST PLUS pipetting robot. For plates which belong to the day 4 to 10 treatment-window, 20µl pure DMEM+++ was added instead. At day 4, medium was removed with the help of a BioTek EL406 washer/dispenser robot and 70 µl of *Balance* medium (including ERBB-inhibitors for treatment windows day 4 to 10, and day 0 to 10 in aforementioned concentrations or excluding ERBB-inhibitors for the day 0 to 4 treatment window). Similarly, wells were refreshed at day 7 with 70 µl *Balance* medium before plates were fixed and immunostained at the experimental endpoint of day 10. After segmentation, only the highest viable concentration was used for further analysis, which were: Cetuximab 3.5 nM, Pertuzumab 25 µg/ml, Lumretuzumab 10 µg/ml, Cetuximab 0.035 nM + Pertuzumab 0.25 µg/ml, and Pertuzumab 25 µg/ml + Lumretuzumab 10 µg/ml.

### scRNA-seq time course

Mature organoids were trypsinized and seeded as described for the maintenance cultures into 6-well culture plates. At 24, 48, 72, 96, 120, 144, 168, 192 and 240 h after seeding, organoids were collected using ice-cold DMEM+++ and singularized as described above. Dissociated cells were filtered through a 40 µm filter (Falcon) and sorted/counted on a BD FACSAria III with a 100 µm nozzle into a 1.5 ml tube. Single viable cells were sorted based on DRAQ7 (1:200) staining and cell size. Post-FACS survival assays after incubation on ice for 30 min were independently performed to ensure cells are viable throughout the whole procedure. We collected via FACS until 12,000 cells per timepoint were collected. For the following timepoints, 12,000 cells could not be reached due to low recovery efficiency: 24 h: 10,286, 48 h: 5,400, 192 h: 8,900. Collected samples were then subsequently used for scRNA-seq library preparation and sequencing.

### scRNA-seq library preparation and sequencing

scRNA-seq library preparation and sequencing was done as detailed elsewhere^5^. In short, libraries were prepared using the GemCode Single Cell 3’ Gel Bead and Library Kit according to the user manual. GEMRT was performed in a Bio-Rad PTC-200 Thermal Cycler with semi-skirted 96-Well Plate (Eppendorf P/N 0030 128.605): 53 °C for 45 min, 85 °C for 5 min; held at 4 °C. After RT, GEMs were broken, and the single strand cDNA was cleaned up with DynaBeads® MyOneTM Silane Beads (Life Technologies P/N 37002D). cDNA was amplified using a Bio-Rad PTC-200 Thermal cycler with 0.2 ml 8-strip nonFlex PCR tubes, with flat Caps (STARLAB P/N I1402-3700): 98 °C for 3 min; cycled 12x: 98 °C for 15 s, 67 °C for 20 s, and 72 °C for 1 min; 72 °C for 1 min; held at 4 °C. Amplified cDNA product was cleaned up with the SPRIselect Reagent Kit (0.6X SPRI). Indexed sequencing libraries were constructed using the reagents in the Chromium Single Cell 3’ Reagents V3.1 (10x Genomics P/N120237), according to the manufacturer’s protocol, with 11 PCR cycles for cDNA amplification. Final libraries were sequenced on a NextSeq500 Illumina platform, with a 56bp cDNA read length. Fastq files were generated using 10X genomics cellranger.

### Long term light sheet imaging

Light sheet imaging was carried out in accordance with a previously described protocol^82^ on a LS1-Live dual illumination and inverted detection microscope from Viventis Microscopy Sàrl was used. Cells were collected from trypsinized mature organoids and seeded into a light sheet holder in 5 µl droplets at a density of 35 cells / µl in a 66% Matrigel + 34% *Start* medium mixture. After a 20 min solidification period, droplets were overlaid with warm *Start* medium. For imaging of the H2B-iRFP670 reporter line, light sheet holders were placed into a 37 °C / 5% CO^2^ incubator for 24 h before transferring the sample onto the light sheet microscope to increase the chance of focusing a surviving cell at the start of the acquisition. On the light sheet microscope, up to 25 organoids were focused on the start of the acquisition and followed over up to 250 h in 15 min (H2B-iRFP670 line) or 30 min (OLFM4-P2A-mNeon-2xNLS line) intervals with a z-stack spanning the whole organoid in 2 µm slicing-intervals. Laser intensity was kept to a minimum necessary to still obtain reasonable signal-to-noise from the raw data, while keeping phototoxicity to a minimum possible. Medium was changed to *Balance* (Table 3) between acquisition intervals at ∼4 days and refreshed at ∼day 7.

### Immunostaining of *in vivo* tissue slices

Slices of normal colonic tissue from anonymous patients were obtained as described (Patients HUB-HS-02-A2-M21-00081 and HUB-HS-02-A2-M21-00225). Samples were formalin fixed and stored in 70% ethanol at 4 °C prior to paraffin embedding (Medite TPC15). Biopsies were subsequently microtome sectioned to a thickness of 3 µm and mounted onto poly-L-lysine coated Poly-Prep glass slides (Cat. No. P0425-72EA, Sigma Aldrich). After incubating the slides at 37 °C over night deparaffination was performed. To this end, the slides were processed in the following steps: 60° for 45 min, 2 × 3 min Neo-Clear (Cat. No. 1.09843, Merck), 2 × 3 min 100% ethanol, 1 × 3 min 96% ethanol, 1 × 70% ethanol, 1 × 5 min ddH2O. Citrate buffer (1 mM Citrate, 0.05% Tween 20, pH 6.0) based antigen retrieval was performed at 95°C for 50 min in a microwave (Milestone Micromed T/T Mega). After cooling down to room temperature (RT) the slides were submerged in blocking buffer (3% donkey serum, 0.5% Triton X in PBS) and incubated for 1 hour at RT. Tissue sections were encircled with a PAP pen and then incubated with primary antibodies: anti-OLFM4 1:200; anti-MUC2 1:200; anti- FABP1 1:200; anti-KI67 1:500; anti-Serotonin 1:1000, anti-Glucagon 1:200 diluted in staining solution (3% donkey serum, 0.1% Triton X in PBS) over night at 4°C. After submerging the slides in PBS for 3 x 20min at RT, sections were stained with secondary antibodies (1:500) and DAPI (1:1000) for 3 hours at RT. After washing in PBS (3 x 20 min, at RT) one drop of mounting medium (Cat. No. 50001, Ibdi) was added before covering the sections with coverslips (0.17 mm).

Tissue sections were imaged with a widefield microscope (Zeiss Axioscan Z1 automated slide scanner) using a Plan Apochromat 20x/0.8 air objective (Zeiss). Fluorophores were excited with a Xcite Xylis LED fluorescence light source (430/475/545/650). The wavelengths were filtered with Ex: BP 470/40; FT 495; Em: BP 525/50 (for Alexa 488), Ex: BP 545/25; FT 570; Em: BP 605/70 (for Alexa 568) and Ex: BP 640/30; FT 660; Em: BP 690/50 (for Alexa 647). Images were collected sequentially with an ORCA flash 4.0V (2048x2048, sCMOS, Hamamatsu) camera. The pixel size was 6.5 µm. 10 planes were collected with a step-size of 0.5 µm at each position. The data was acquired as z-series first, then channel, and position. The z-slice with the highest sharpness was selected to be displayed in the figures.

### Reporter line creation

For video S1 and S2, **a** human colon line (HUB-02-A2-040) was lentivirally transduced with pGK Dest H2B-miRFP670 (Addgene). Lentiviral transduction was performed on single cells after 0.05% Trypsin EDTA (Gibco) dissociation. Monoclonal organoids were produced by isolating and sorting positive miRFP cells using a FACS-ARIA III (BD Biosciences) after two rounds of passages.

For the light-sheet movies following stem cell dynamics (Video S3-S5), we used the human colon line (HUB-02-A2-040). The sgRNA was selected based on the Benchling website (https://benchling.com) using their CRISPR-tool (taking on-off target selection into account^83^). The gRNA overlaps with the stop codon (Table S1). The target sequence was ordered as fwd and rev oligos (Microsynth) and cloned into a generated minimalistic targeting backbone miniBackBone (2.5kb – hU6-sgRNA) following the protocol described before^84^

Human colonic organoids were transiently transfected using a NEPA21 electroporator and a previously developed protocol^85^. For electroporation, organoids were dissociated into single cells and washed twice with OptiMEM. The resulting pellet was resuspended in BTXpress solution (BTX) with 15 μg of plasmids, after which electroporation was performed.

For the generation of CRISPR-HOT reporter organoids (OLFM4-P2A-mNeon-2xNLS-P2A- blast.) see previous description^86^. 4 days after electroporation and culture in *Start* medium, we started selecting in our *Balance* medium with blasticidin (5 µg/ml) and picked single organoid clones. From the single clones mNeongreen positive cells were sorted using a FACS-ARIA III (BD Biosciences).

In short, we used a targeting plasmid containing a fluorescent protein that localizes to the nucleus and selection cassette (P2A-mNeongreen-2xNLS-P2A-bast.) that is linearized at a defined base position by a specific sgRNA and Cas9 provided from a second plasmid, here one could pick one out of three frame-selector plasmids, which also encode mCherry^87^. These two plasmids together with the plasmid encoding the sgRNA for the locus of interest are co-electroporated (**Table S1**).

### Fluorescence-activated Cell Sorting (FACS) and analysis

On Day 10, organoids were singularized, resuspended in PBS, filtered through a 40- micrometer filter, and counted. DAPI was applied for 5 minutes to detect dead cells, followed by fixation with 4% PFA for 15 minutes. The cells were then resuspended in PBS at a concentration of 2*10^5 cells per mL for antibody staining. Permeabilization was carried out using a solution containing 0.5% Triton X-100, 100 mM NH4Cl, and 3% donkey serum in PBS for 15 minutes at room temperature. Prior to staining, a permeabilization step was carried out in 0.5% Triton X-100, 100 mM NH4Cl, 3% donkey serum in PBS solution for 15 min at RT.

Primary antibodies (i.e., MUC2, FABP1, OLFM4), at the same dilutions as previously described for IF staining of organoids, were applied and incubated for 10 minutes, followed by centrifugation at 900g for 5 minutes and removal of the supernatant. The cells were washed three times with PBS and then left overnight at 4 °C. Subsequently, samples were then overlayed with secondary antibodies against the used primary species (1:500). Negative controls were included by staining only with secondary antibodies. After a final wash with PBS, the cells were resuspended in 200 µl of PBS for FACS measurement on the MA900 cell sorter (SONY).

FACS strategies for the reporter lines were performed on the FACS-ARIA III (BD Biosciences) (Supplementary information 1).

### Image analysis

All image analysis steps were conducted in Python (version 3.11.3) unless stated otherwise.

#### Raw-image preprocessing

Raw-images were first, if not done by the microscope-software during the data acquisition, converted into maximum-intensity-projections (MIPs). Individual MIPs of single field-of-views (FoVs) were then stitched together based on their central coordinates within one well to generate whole-well MIPs.

#### Organoid-level segmentation

Whole-well MIPs of the DAPI-channel were segmented using different neural-network-based segmentation workflows based on the fixation timepoint after single cell seeding. Samples fixed and imaged 2 days after seeding of single cells were segmentation with a custom model using RDCNet (version 0.1,^88^). Samples imaged at all other timepoints after single cell seeding were segmented in CellPose^89,90^ (version 2.2, run in Python 3.8.17) using multiple custom-trained models modified from the cyto2 model. For samples fixed after day 4, MIPs were temporarily binned with a factor of 4 in x and y direction to improve segmentation speed. Thereafter, segmentation masks were upscaled by the same factor to fit the size of native MIPs again. Furthermore, a median filter was used on segmentation masks to reduce binning-associated roughness of edges. Before further image processing, such as feature extraction and quality control, regions of individual objects were extracted from the MIPs of all acquired wavelengths and saved individually using the generated segmentation mask and ID of the object in combination with a dilated bounding box. All segmentation models are available upon request.

#### Nuclei-level segmentation

Nuclei from day 10 old human colonic organoids were segmented using the in-built neural-network (nuclei) from CellPose^89^. To convert the anisotropic into isotropic data for visualization purposes we used in Python (version 3.9.15) the scikit-image tool to rescale. Confocal imaging of fixed organoid samples was performed using the Nikon Ti2-E Eclipse Inverted motorized stand with the Yokogawa CSU W1 with Dual camera (CAM1, X-11424; CAM2, 11736) T2 spinning-disk confocal scanning unit, and Plan-Apo ×25/1.05 NA silicon-immersion objective. The laser lines used included Toptica iBeam Smart 405/488/639 nm and Cobolt Jive 561 nm. Image stacks were acquired with a slice thickness of 1 μm. All MUC2 positive cells were counted manually.

#### Feature extraction

Feature extraction was done on region-extracted object MIPs. First, intensity measurements of all acquired wavelengths of an object were done within the segmentation mask of the individual object to exclude object-external signal. Channel-imaged were blurred with a Gaussian kernel (sigma = 3) and the mean, minimum, maximum, standard-deviation as well as quantiles (0.25, 0.50, 0.75, 0.99) and weighted image-moments (from the scikit-image regionprops function, version 0.20.0) were calculated. The pixel-wise correlation between different staining markers within one object (excluding DAPI-mediated DNA-staining) was calculated using the corrcoef function of NumPy (version 1.24.3). Furthermore, intensity measurements were conducted in the area defined as positive for a staining. For this, thresholds have been initially set automatically by probing the threshold found in 20 randomly chose objects per timepoint per utilized staining using the triangle thresholding method from the scikit-image package on gaussian blurred (sigma = 3) images. The mean of all these thresholds was tested visually on randomly chosen organoids of all timepoints (in the case of multiple probed sampling timepoints) and adjusted if necessary. The found threshold was then imposed onto the channel image of all objects and aforementioned intensity features were calculated within the thresholded area. Furthermore, the absolute size of the positive area as well its relative size compared to the total size of the organoid was extracted. Finally, the asymmetry between the centroids of the segmentation mask and thresholded area was calculated. Next, morphological features were calculated on the segmentation mask using the regionprops function of scikit-image. In addition, the circularity of the object, its ratio between the major and minor axis, and the number of cavities were calculated. For features describing the number and morphology of crypt-structures, the segmentation mask was initially skeletonized using the morphology package of scikit-image (method = ‘lee’, sigma = 3). Additionally, a maximum-inscribed circle with half of its radius was fitted onto the organoid mask using code from https://gist.github.com/DIYer22. Next, skeleton branches within the circle were deleted and endpoints of remaining branches elongated until the segmentation border using the same average directionality as the last 50 pixels of that branch. Intermediate branches which did not touch the segmentation border (i.e., branches start at the maximum- inscribed circle and further bifurcate into multiple other branches) were deleted and remaining branches were counted, as well as their total length summed-up. Finally, the bounding box of organoids was calculated and shrunk by 1 pixel in all directions. Next, the longest straight stretch of overlay between the organoid outline and bounding box was calculated as prolonged straight lines indicate not fully imaged objects at the imaging-border. Features were minmax normalized between 0 and 1 using the preprocessing function of scikit-learn (version 1.2.2). Normal distribution of features was analyzed with a quantile-quantile plot and the resulting square root of the coefficient of determination (probplot function of scipy, version 1.10.1). Features which were not normally distributed were largely associated with area measurements, thus all area-associated measurements were transformed using the natural logarithm of the input array +1. Finally, features were saved into an AnnData (version 0.9.1) object.

#### Quality control

Before further processing, resulting objects were filtered on a subset of their features. Specifically, organoids with a straight-border fraction of >= 0.4, a low DAPI mean intensity, and an area smaller than a single cell were excluded from further analysis. Remaining objects were visually probed, and filters adjusted if needed, until >99% of objects were correctly segmented organoids. Finally, plate-position bias was checked by comparing average areas, circularities, and outgrowth efficiencies of wells within one condition.

#### Dimension reduction, phenotyping and trajectory analysis

For trajectory analysis, all organoids were initially placed into a UMAP (umap-learn package, version 0.5.3) based on all morphological organoid-level minmax normalized (and, in the case of area-associated features, log1p transformed) features. The UMAP parameters were set as follows: NRG1/EGF titration experiment: n_neighbors = 50, min_dist = 0.4; time course experiments: n_neighbors = 25, min_dist = 0.4; ERBB inhibition experiment: n_neighbors = 100, min_dist = 0.3. All other parameters used default values. Next, grouping into clusters with high morphological similarity was conducted using PhenoGraph (version 1.5.7) using experiment-specific k-values (NRG1/EGF titration experiment: k = 20; time course experiments: k = 70; ERBB inhibition experiment: k = 30) and default values for all other parameters. Next, clusters were assigned to phenotypes based on their morphology. For the ERBB inhibition experiments, we found a cluster largely consisting of organoids with segmentation artefacts. This cluster has been removed and the dimension reduction with UMAP, clustering by PhenoGraph, and cluster- naming has been re-run. Next, pseudotime trajectories were computed using the SlingShot package in python (pySlingshot, version 0.0.2) with default parameters without defining end-states. Start_node was set as the cluster with the smallest average organoid size. The fit was run three times and the average trajectory and pseudotime was saved for further analysis. Pseudotimer numbers were minmax normalized between 0 and 1 for all trajectories and binned into 10 bins of equal size (each spanning a pseudotime frame of 0.1).

### scRNA-seq analysis

All analyses, unless stated otherwise, were performed within R (version 4.1.3,^91^) and Bioconductor (version 3.14, ^92^).

#### Alignment and quantification

The count table for the human colon organoid dataset was created using the Cell Ranger software (version 4.0.0) with default parameters and the GRCh38 human genome assembly as a reference. To count intronic as well as exonic reads separately, we created a custom gene annotation gtf file with exonic as well as intronic annotation. We downloaded the RefSeq transcript annotation file ncbiRefSeq.txt from UCSC (date: 17.09.21) with its corresponding Entrez gene id mapping file ncbiRefSeqLink.txt. We first removed transcripts that were not assigned (chrUn), assigned to random (random), alt (alt), or fix (fix) chromosomes. Secondly, we removed transcripts that were assigned to multiple loci in the genome. Finally, we removed genes that were assigned to multiple chromosomes. For each gene, we determined the gene body coordinates by calculating the minimum start and maximum end coordinates of all exons assigned to a particular gene. To obtain the intronic coordinates for every gene, we determined all the regions in each gene body that were not covered by exons using the setdiff operation from the R package GenomicRanges. To avoid spurious intronic assignments close to exon junctions, we extended the exon coordinates by 10bp before performing the setdiff operation. We combined the exonic and intronic coordinate tables into a single one, adding a suffix of “_EX” for gene and transcript identifiers corresponding to exonic regions and a prefix of “_IN” for regions corresponding to intronic regions. We exported that table as a gtf file using the function export from the R package rtracklayer.

#### Quality control, filtering, and normalization

For each cell, the total UMI count and the fraction of UMI counts mapping to mitochondrial genes were calculated. High quality cells, defined as cells with a mitochondrial fraction less than X (89th percentile) and a total UMI count greater than 4500, were retained for downstream analysis. Low expressed genes, defined as genes that were detected in less than 27 cells (0.05% of all cells), were removed, resulting in a final count matrix of 18,630 genes by 53,640 cells. Raw UMI counts were normalized and log2-transformed using *log2(nig · si + 1)*, where nig is the raw UMI count for gene *g* in cell *i*, and *si* is the size factor for cell *i* calculated as *si = mean(N)/Ni*, where *mean(N)* is the arithmetic mean of the sums of UMI counts for each cell, and *Ni* is the sum of UMI counts for cell *i*. Unless otherwise specified, log2-normalized counts were used in all downstream analyses.

#### Identification of highly variable genes

Highly variable genes for use in dimensionality reduction were identified from the pool of all cells using the *modelGeneVar* and *getTopHVGs* functions with arguments *var.threshold = 0* and *fdr.threshold = 0.05* from the *scran* package (version 1.22.1, ^93^).

#### Cell cycle correction

We calculated an initial embedding of the cells using *runPCA* on variable genes and *runUMAP* on the PCA result from the *scater* package (version 1.22.0, ^94^), which predominantly separated cells by their cell cycle phase, predicted using the *tricycle* package (version 1.2.1, ^95^) to estimate cell cycle position and assigned to phases with the following intervals: G1 [0, 0.8) or [5, 2ρχ], S [0.8, 2.8), G2M [2.8, 5). Next, we used the *getVarianceExplained* function from *scater* (version 1.22.0, ^94^) to identify genes explaining more than 10% of the predicted cell cycle phases. A newly calculated PCA and UMAP embedding excluding these genes showed mixing of cells from different cell cycle phases.

#### Sketching and dimensionality reduction

An initial dimensionality reduction was calculated by running *calculatePCA* with default parameters from scater (version 1.22.0, ^94^), excluding cell cycle associated genes identified above. In order to reduce bias towards highly abundant cell types, we identified a representative subset of cells using the geosketch algorithm^42^ from the *sketchR* package (version 0.1.4, ^96^), with the initial PCA as input and selecting 6,437 cells (12% of all cells). These sketched cells were used to identify highly variable genes and calculate a PCA embedding using *runPCA* from *scater* (version 1.22.0, ^94^). Non-sketched cells were projected accordingly using the obtained rotation matrix and scaling parameters, thereby generating an embedding of all cells in the same space (PCAsketch). Similarly, a UMAP embedding for all cells was created by first calculating a UMAP model using *umap* from the *uwot* package (version 0.1.11, ^97^) with the PCA from sketched cells as input and parameters *n_components = 2*, *min_dist = 0.4* and *n_neighbors = 10*, and then using this model to project all cells (using PCAsketch as input) into the same UMAP space (UMAPsketch). Compared to embedding derived from all cells, the sketched embeddings provide a better separation of rare, related cell types (for example EC and EE non**-**EC).

#### Cell type annotation

We first clustered the sketched cells by constructing a nearest neighbor graph with *FindNeighbors* and running *FindClusters* from Seurat (version 4.1.0, ^98^) with *resolution = 6.5.* Cluster annotations were transferred to the non-sketched cells by k nearest neighbor matching using the *knn* function from the package *class* (version 7.3-20, ^99^) with PCAsketch to calculate cell distances and *k = 1*. The resulting 46 clusters were manually mapped to known cell types using published marker genes, yielding 20 distinct cell labels.

#### Diffusion maps

Diffusion maps were calculated using *DiffusionMap* from *destiny* (version 3.8.1, ^100^) on PCAsketch and with *sigma = 9.8* (identified by *find_sigmas* and *optimal_sigma*) and *knn = 100* (identified by *find_dm_k*).

#### Pseudotime estimation

Pseudotime was estimated on diffusion maps using the *floodPseudotimeCalc* function from *URD* (version 1.1.1, ^101^) with default parameters and stem cell progenitors as starting population. 150 iterations (calls to *floodPseudotimeCalc*) were run for each set of starting cells and processed by *floodPseudotimeProcess* with *stability.div = 20*. As the starting cells will get a pseudotime of zero by definition, we modified the published workflow by repeatedly splitting the set of starting cells randomly into two equally sized subsets and running the pseudotime estimation separately on each half, thereby assigning non-zero pseudotime values to the other half. This procedure was repeated eight times, resulting in a total of 16 pseudotime estimates for all non-starting cells, and 8 estimates for all starting cells. The final pseudotime for each cell was calculated by averaging the individual run results. The pseudotime estimates for dedifferentiating cell types (“Colonocyte reprogramming”, “Secretory reprogramming”, “Stem cells reprogramming”, “YAP+CLU”) were multiplied with -1, to account for the fact that these cells are changing from mature to undifferentiated states. To smooth the transition around zero, we subtracted from the set of negative pseudotime values the 99^th^ percentile, and the first percentile from the positive set. This procedure allows to describe the system with a single pseudotime axis, where negative and positive values correspond de- and redifferentiating cells, respectively.

#### Gene expression over pseudotime

For visualization of gene expression as a function of pseudotime, sketched cells belonging to a given lineage (as indicated in the visualization) were selected and the log2-normalized expression values were smoothed over pseudotime using the *loess* function from R (version 4.1.3) with *span = 0.9.* The resulting fit was used to predict smoothed expression values with standard errors (parameter *se = TRUE*), and finally the smoothed expression curve with standard errors was scaled into the interval [0,1] for plotting.

#### Similarity of mature cell types to published datasets

In order to calculate the similarity of mature cell types to annotated cell types in published datasets, we first manually mapped the provided cell labels to the four major intestinal cell lineages (“Stem cells”, “Goblet cells”, “Enteroendocrine cells”, “Colonocytes”). Pseudo-bulk expression profiles were calculated for both own and public data set per major cell lineage, using *aggregateAcrossCells* from the *scuttle* package (version 1.4.0, ^94^) with *use.assay.type = “logcounts”* and *statistics = “mean”*. A common set of genes was identified by intersecting genes detected in all datasets and available sets of highly variable genes (own analysis and data from^12^), resulting in 354 genes. Pairwise Pearson correlation coefficients were then calculated between all pairs of own and published cell lineage pseudo-bulks and used for visualization.

#### Integration of public and organoid single cell data

In order to integrate own and published single cell data into the same reduced dimensional space, we used raw (unnormalized) counts where available (own, ^12,66^). For datasets where only log-normalized values were available, we reverse-transformed these to counts by exponentiating, subtracting the pseudocount and then dividing by the scaling factors described in the source publications^65^. We then used *SCTransform* from *Seurat* (version 4.3.0, ^98^) with *method = “glmGamPoi”* to re-normalize all data in a consistent way. Data sets were integrated using *Seurat* following the package documentation: We first called *SelectIntegrationFeatures* with *nfeatures = 3000*, then *PrepSCTIntegration*, followed by *FindIntegrationAnchors* with *normalization.method = “SCT”*. Finally, we used these intermediate results in *IntegrateData* with *normalization.method = “SCT”*. The integrated data was embedded into reduced dimensional space for visualization using *Seurat*’s *RunPCA* and *RunUMAP* with default parameters. Cell types were then re-annotated on the integrated data similarly as described above, with *FindNeighbors* and *FindClusters* from Seurat (version 4.3.0, ^98^) with *resolution = 2.5.* The resulting 71 clusters were manually mapped to known cell types using published marker genes, yielding 14 distinct cell labels.

#### Clustering of datasets by cell type frequencies

Cell type frequencies for the integrated data were calculated per individual patients (*in vivo* data) or per culture medium (organoids), and clustered using R’s *hclust* function with *method = “ward.D2”* and Euclidean distances.

