## Supplementary Information for "Dynamics and plasticity of stem cells in the regenerating human colonic epithelium"

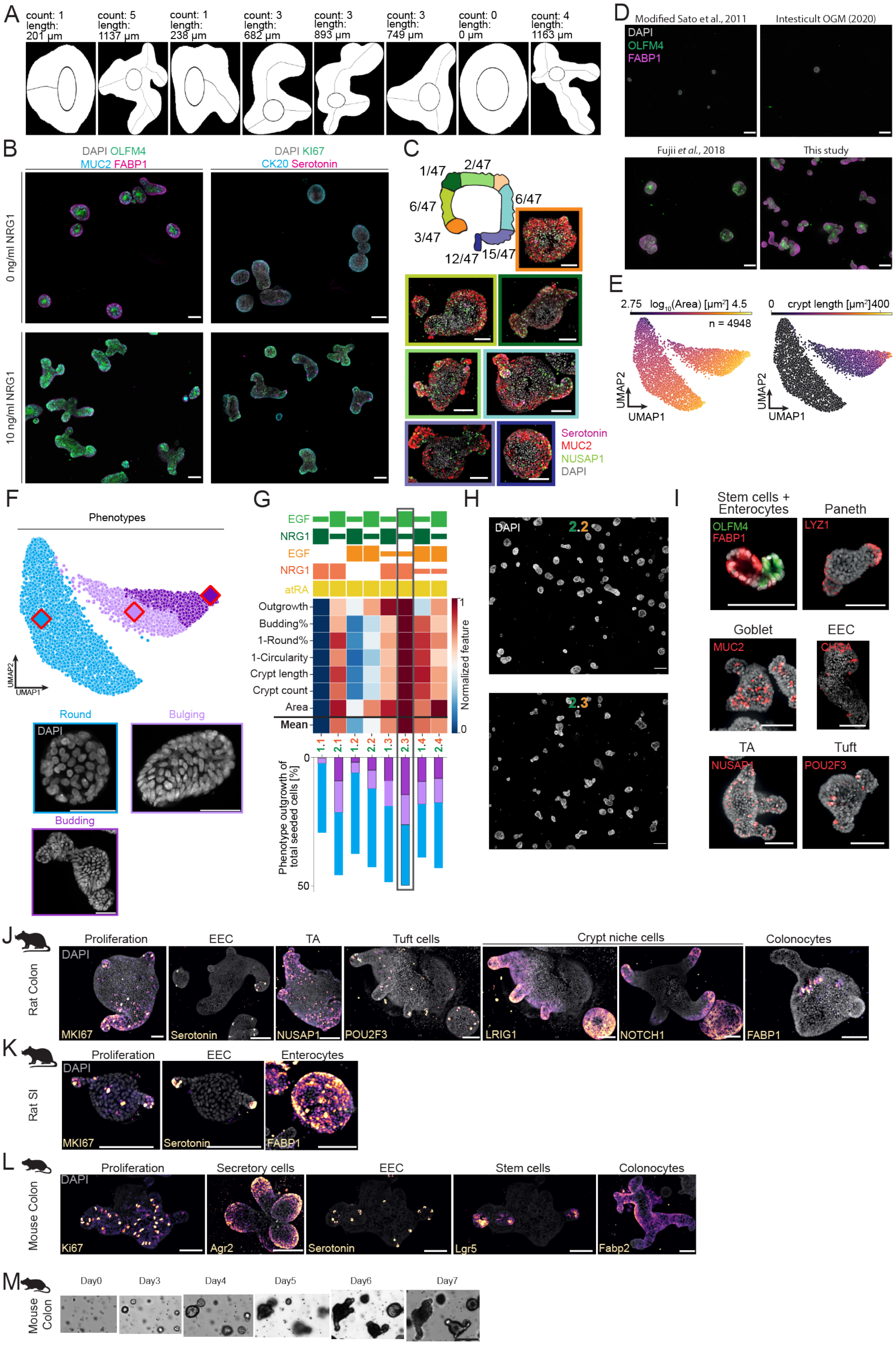

A
count: 1 length: 201 µm
count: 5 length: 1137 µm
count: 1 length: 238 µm
count: 3 length: 682 µm
count: 3 length: 893 µm
count: 3 length: 749 µm
count: 0 length: 0 µm
count: 4 length: 1163 µm
B
DAPI OLFM4 MUC2 FABP1
DAPI KI67 CK20 Serotonin
0 ng/ml NRG1
10 ng/ml NRG1
C
1/47
2/47
6/47
6/47
3/47
12/47
15/47
Serotonin
MUC2
NUSAP1
DAPI
D
Modified Sato et al., 2011
Intesticult OGM (2020)
DAPI
OLFM4
FABP1
Fujii et al., 2018
This study
E
2.75 log10(Area) [µm²] 4.5
n = 4948
0 crypt length [µm²] 400
UMAP2
UMAP1
F
Phenotypes
Round
Bulging
Budding
DAPI
G
EGF
NRG1
EGF
NRG1
atRA
Outgrowth
Budding%
1-Round%
1-Circularity
Crypt length
Crypt count
Area
Mean
Normalized feature
1.1 2.1 1.2 2.2 1.3 2.3 1.4 2.4
Phenotype outgrowth of total seeded cells [%]
0
50
H
DAPI
2.2
2.3
I
Stem cells + Enterocytes
OLFM4
FABP1
Paneth
LYZ1
Goblet
MUC2
EEC
CHGA
TA
NUSAP1
Tuft
POU2F3
J
Rat Colon
Proliferation
EEC
TA
Tuft cells
Crypt niche cells
Colonocytes
DAPI
MKI67
Serotonin
NUSAP1
POU2F3
LRIG1
NOTCH1
FABP1
K
Rat SI
Proliferation
EEC
Enterocytes
DAPI
MKI67
Serotonin
FABP1
L
Mouse Colon
Proliferation
Secretory cells
EEC
Stem cells
Colonocytes
DAPI
Ki67
Agr2
Serotonin
Lgr5
Fabp2
M
Mouse Colon
Day0
Day3
Day4
Day5
Day6
Day7

**Figure S1 | Development of a niche-inspired medium composition through dynamic EGF-like ligand supplementation.** ***Related to Figure 1*** (**A**) Exemplary results of the crypt-morphogenesis analysis workflow. Shown are the segmentation masks of eight randomly picked organoids and quantification of their crypt count and total crypt length (see Methods). (**B**) Representative full-well images of human colonic organoids grown without (upper row) and with (lower row) NRG1 supplementation. Scale bars, 200 µm. (**C**) Exemplary colonic organoids from 47 individuals grown in the developed medium combination (2.3.3) from seven anatomical subsections of the colon. Origin of biopsy is indicated by the image border (orange: ceacum, light green: ascending colon, dark green: hepatic flexure, neon green: transverse colon, cyan: descending colon, light purple: sigmoid colon, dark purple: rectum). Numbers indicate distribution of subsection among all samples. Scale bar, 50 µm. (**D**) Representative full-well images of mature human colonic organoids grown from single cells (125 cells/well) in media modified from Sato et al., 2011 (upper left panel), Fujii et al., 2018 (lower left panel), a commercial option (human Intesticult OGM 2020, upper right panel) and the medium of this study (lower right panel). Scale bar, 200 µm. (**E-I**) Results of time-dependent EGF-like ligand supplementation on human small intestinal organoids. (**E**) Selected features highlighted on a UMAP based on the shape-feature space of mature small intestinal organoids grown in 8 different conditions (see Table 2). (**F**) Three major observed phenotypes ('Round': light blue, 'Bulging': light pink, 'Budding': purple) derived from shape-features and exemplary images of small intestinal organoids within this phenotype. Diamonds with red border show coordinates of the organoids used as phenotype examples. Scale bars, 50 µm. (**G**) Heatmap of row-normalized features associated with successful crypt-morphogenesis among all 8 tested conditions (upper panel, see Table 2) and the outgrowth efficiency of tested conditions (lower panel). Outgrown small intestinal organoids are colored based on their phenotype. Grey rectangle indicates medium combination 2.3, which is used for further experimentation. Size of bars above heatmap represents the concentrations of EGF and NRG1 until day 4 (green), as well as EGF, NRG1 and all-trans retinoic acid (atRA) from day 4 to day 10 (orange). Compared to the medium testing for colonic organoids, no WNT-NGS was used from day 4 onwards. Instead atRA was supplemented to improve enterocyte differentiation. (**H**) Representative whole-well images of a poor (2.2) and well (2.3) performing medium combination. Scale bars, 200 µm. (**I**) Representative images of small intestinal organoids grown in medium combination 2.3 and immunostained for major human small intestinal cell types as well as DNA. Scale bars, 100 µm. (**J-L**) Representative images of organoids derived from rat colonic (**J**), rat small intestinal (**K**), or mouse colonic (**L**) tissue grown in ***(Start/Balance)*** medium combination and immunostained for major intestinal markers as well as DNA. Scale bars, 100 µm. (**M**) Mouse colon organoid maturation from single cells captured by brightfield imaging. Scale bars, 100 µm. All images are maximum intensity projections (MIPs) of z-stacks acquired on a spinning-disk confocal system.

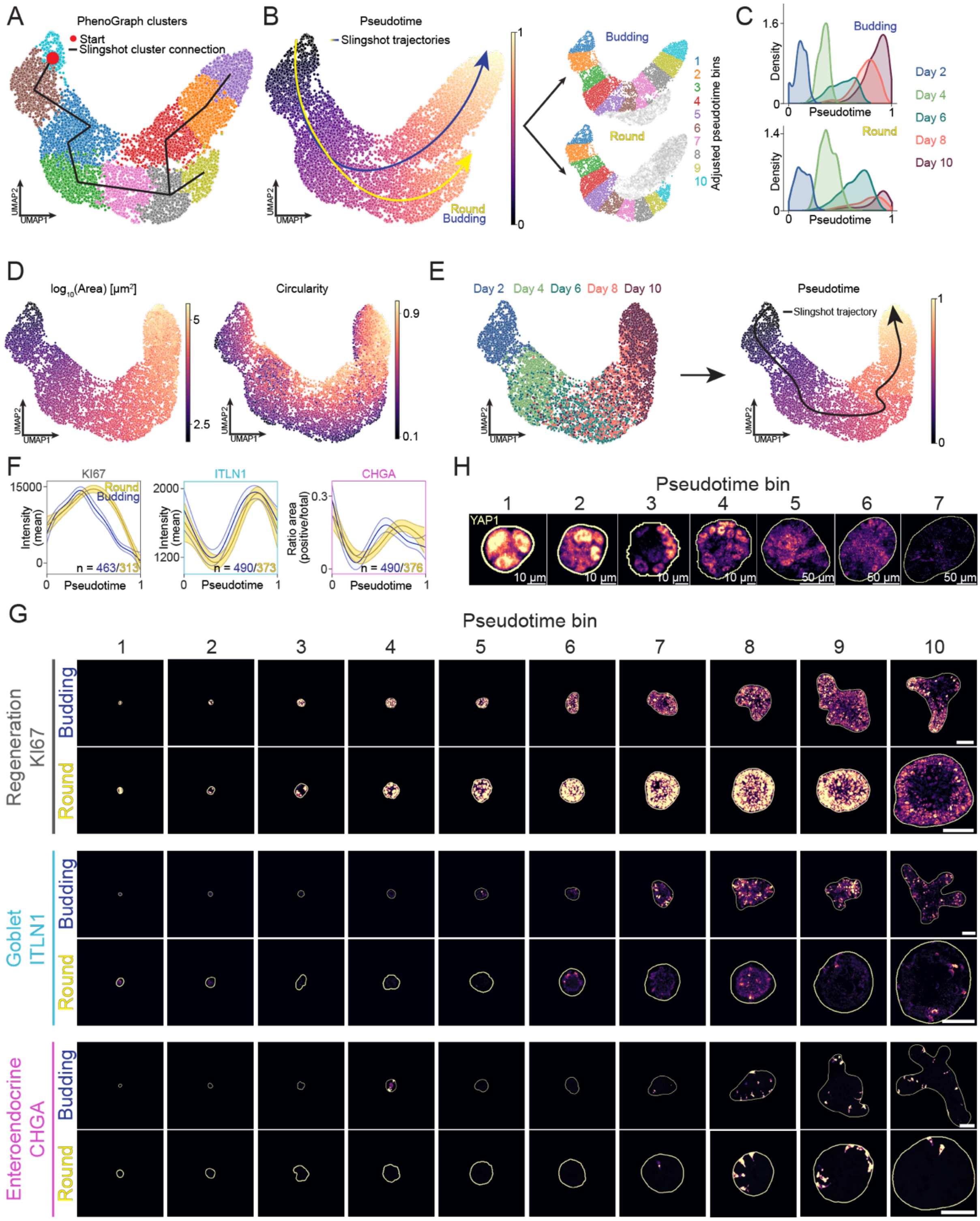


**Figure S2 | Image-based pseudotime analysis indicates two distinct maturation trajectories.** ***Related to Figure 2*** (**A-C**) Results of the Slingshot algorithm. (**A**) PhenoGraph clusters shown on the shape-feature UMAP with the start cluster (red dot) and found connections by Slingshot. (**B**) Resulting pseudotime from the Slingshot algorithm including identified trajectories as well as the 10 derived pseudotime bins for both trajectories. (**C**) Organoid-density plots over pseudotime by fixation timepoint for both organoids in the 'Budding' (top) or 'Round' (bottom) trajectory. (**D**) Organoids from time course experiments placed in an UMAP based on their shape-feature space without crypt-associated features (crypt count and total crypt length). Highlighted are their area (left) and circularity (right). (**E**) UMAP based on the shape-feature space without crypt-associated features (crypt count and total crypt length) highlighting organoids from the five fixation timepoints and their resulting pseudotime. Without crypt-associated features, Slingshot fails to detect multiple maturation trajectories. (**F**) Predicted mean intensity by a generalized additive model (GAM) including 95% confidence intervals of markers associated with proliferation and cell type identity over pseudotime. (**G**) Representative images of marker expression in organoids along detected pseudotime bins in both trajectories. White outline represents the segmentation mask. Scale bars, 100 µm. (**H**) Zoom-in of YAP1 expression showing nuclear translocation in representative organoids between pseudotime bin 1 and 7 of organoids in the 'Budding' trajectory. Scale bars, 10 µm. All images are maximum intensity projections (MIPs) of z-stacks acquired on a spinning-disk confocal system.

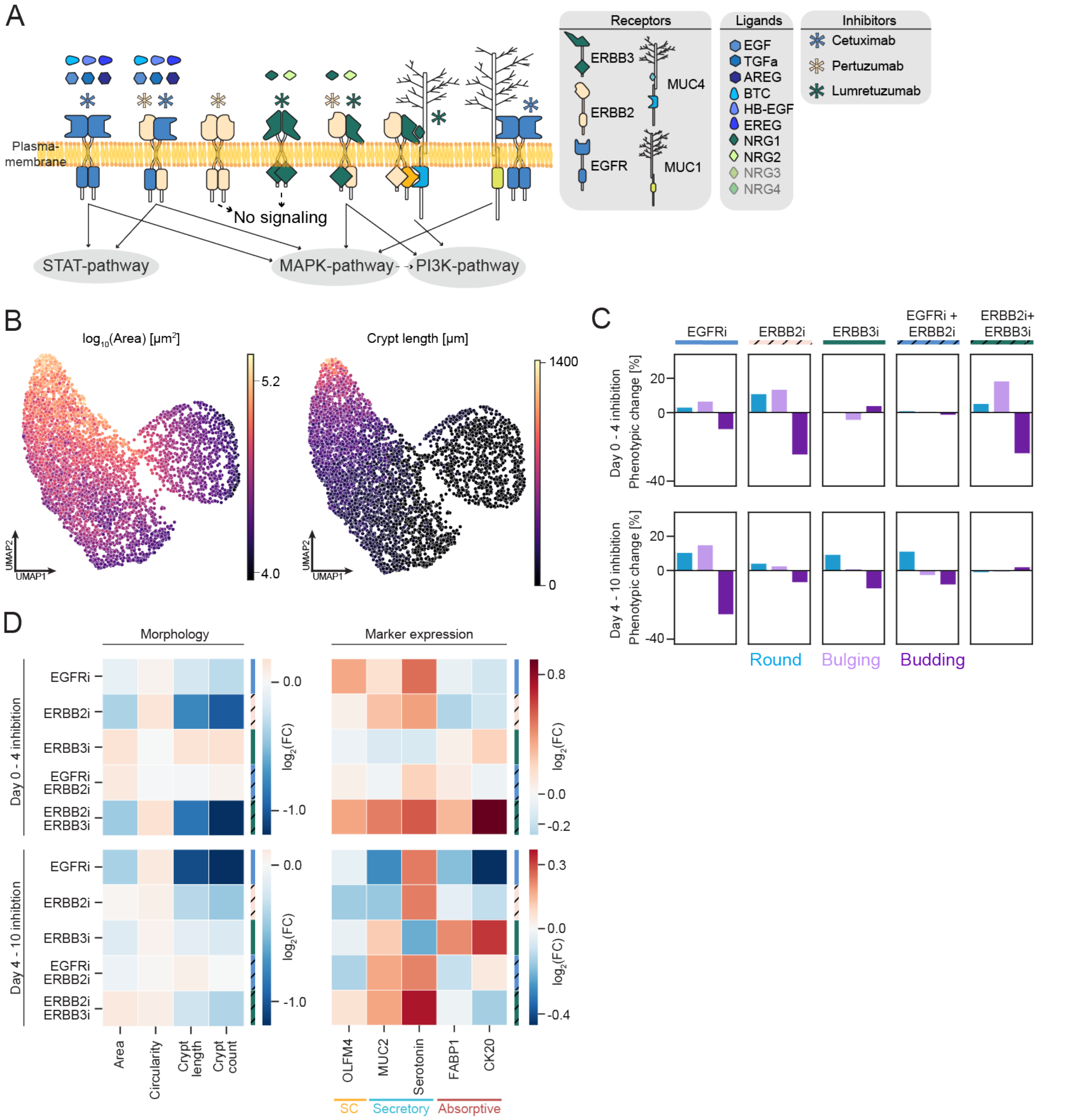


**Figure S3 | Differential ERBB-receptor activation influences lineage specification in human colonic organoids.** ***Related to Figure 3*** (**A**) Extended, schematic representation of the ERBB-receptor class signaling in the intestinal epithelium and the experimental approach. Single cells derived from mature organoids were seeded at day 0 and grown for 10 days. During this time, ERBB receptors (EGFR, ERRB2, and ERBB3) were inhibited by antibody treatments either as a single inhibition, or in combination with ERBB2 inhibitor to prevent heterodimer formation. The inhibition was conducted in two time windows (day 0 to 4 and day 4 to 10). (**B**) Organoids from the ERBB-receptor inhibition (pooling all treatment windows and concentrations) placed in an UMAP based on their shape-feature space color-coded by their area (left), total crypt length (right). (**C**) Change in phenotypic composition of ERBB-inhibition experiments during the two tested time-windows (rows) relative to the corresponding control. The color of the bar indicates the phenotype (‘Round’: light blue, ‘Bulging’: light pink, ‘Budding’: purple). (**D**) Heatmap showing the $\log_2$(fold change) of organoid-level morphological (left) and marker expression (mean intensity) features of used ERBB-receptor inhibition treatments compared to the corresponding control.

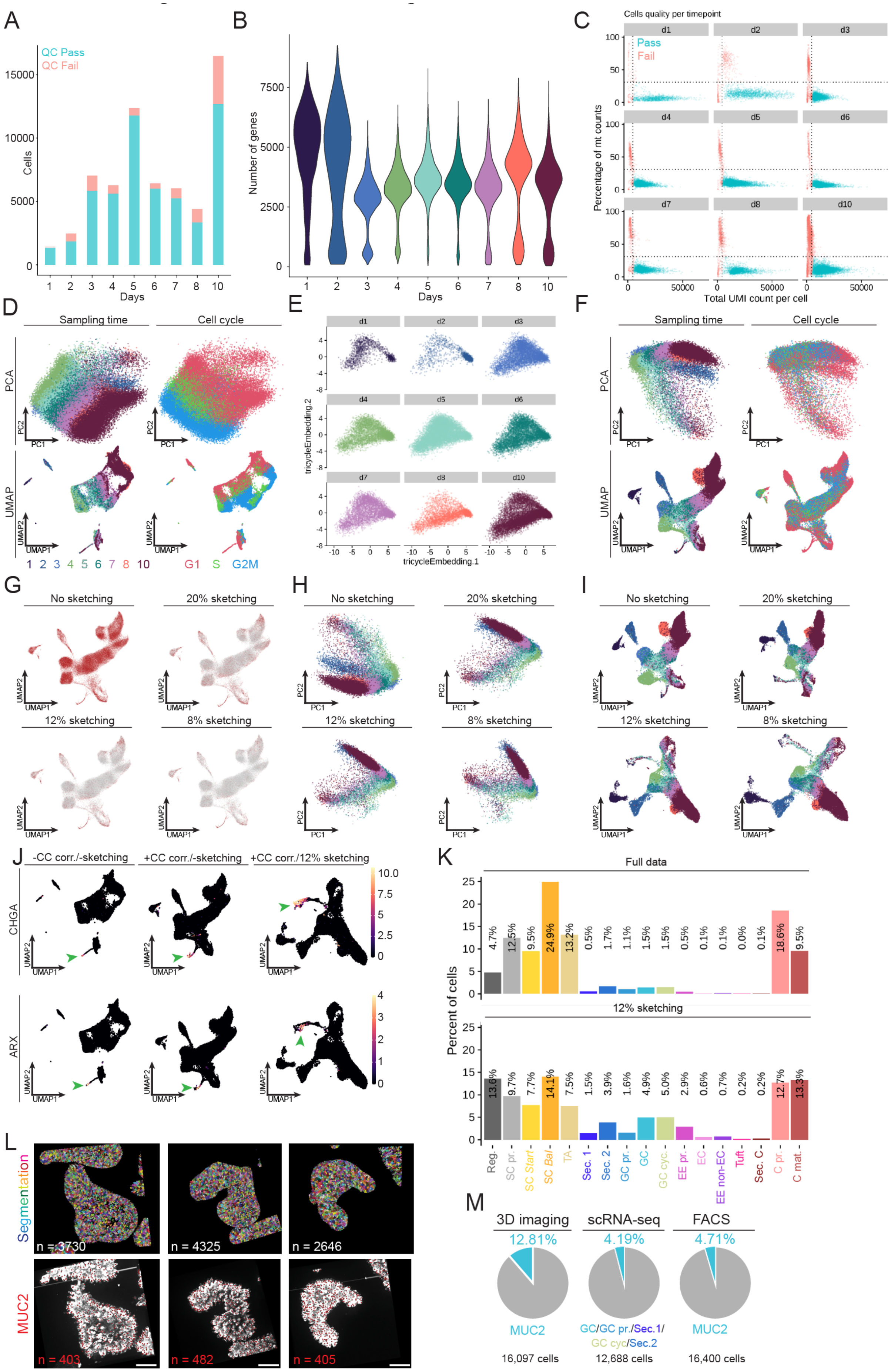
A
QC Pass
QC Fail
Cells
Days
B
Number of genes
Days
C
Cells quality per timepoint
Pass
Fail
Percentage of mt counts
Total UMI count per cell
d1
d2
d3
d4
d5
d6
d7
d8
d10
D
Sampling time
Cell cycle
PCA
UMAP
1 2 3 4 5 6 7 8 10
G1 S G2M
E
tricycleEmbedding.2
tricycleEmbedding.1
F
Sampling time
Cell cycle
PCA
UMAP
G
No sketching
20% sketching
12% sketching
8% sketching
H
No sketching
20% sketching
12% sketching
8% sketching
I
No sketching
20% sketching
12% sketching
8% sketching
J
-CC corr./-sketching
+CC corr./-sketching
+CC corr./12% sketching
CHGA
ARX
K
Full data
12% sketching
Percent of cells
4.7% 12.5% 9.5% 24.9% 13.2% 0.5% 1.7% 1.1% 1.5% 1.5% 0.5% 0.1% 0.1% 0.0% 0.1% 18.6% 9.5%
13.6% 9.7% 7.7% 14.1% 7.5% 1.5% 3.9% 1.6% 4.9% 5.0% 2.9% 0.6% 0.7% 0.2% 0.2% 12.7% 13.3%
Reg.
SC pr.
SC Start
SC Bal
TA
Sec. 1
Sec. 2
GC pr.
GC
GC cyc.
EE pr.
EC
EE non-EC
Tuft
Sec. C
C pr.
C mat.
L
Segmentation
MUC2
n = 3730
n = 4325
n = 2646
n = 403
n = 482
n = 405
M
3D imaging
scRNA-seq
FACS
12.81%
4.19%
4.71%
MUC2
GC/GC pr./Sec.1/
GC cyc/Sec.2
MUC2
16,097 cells
12,688 cells
16,400 cells

**Figure S4 | scRNA-seq reveals all major cell lineages in the mature organoid.** ***Related to Figure 4*** (**A-C**) Quality control (QC) of the acquired scRNA-seq data. (**A**) Fraction of cells which passed (yellow) and failed (red) the quality control steps over their sampling timepoint. (**B**) Number of detected genes per cell over sampling timepoints. (**C**) Counts of detected unique molecular identifiers (UMI) within a and the fraction of detected mitochondrial genes. Cells with a low number of detected UMI and/or high percentage of mitochondrial genes are removed (red). (**D-F**) Cell-cycle (CC) correction (see Methods). (**D**) First two principal components of a principal component analysis (PCA, upper row) and UMAP (lower row) of the scRNA-seq data color-coded by sampling timepoint (left panels) and cell-cycle (right panel). (**E**) Tricycle-embedding of cells across sampling timepoints. (**F**) First two principal components of a PCA (upper row) and UMAP (lower row) of the scRNA-seq data color-coded by sampling timepoint (left panels) and cell-cycle (right panel) of cell-cycle corrected data. (**G-H**) Cell sketching (see Methods). (**G**) Cells used for low-dimensional embedding (red) among all cells (gray) highlighted in a UMAP of the cell-cycle corrected scRNA-seq data. Three different percentages cells used for sketching are indicated. (**H**) Scatter plot of the first two principal components of a PCA resulting from either the full cell-cycle corrected data (upper left), or varying sketching percentages. (**I**) UMAP of the first two principal components of a PCA resulting from either the full cell-cycle corrected data (upper left), or varying sketching percentages. (**J**) Expression of two rare markers (CHGA (upper row) and ARX (lower row)) in UMAPs of either the native data (left), cell-cycle corrected (middle), or cell-cycle corrected and sketched data (right). Cluster of positive cells is indicated by arrows in magenta. (**K**) Percentage of cells belonging to annotated clusters from the native data (top) and the sketched data (bottom). (**L**) 3D single-cell segmentation of three organoids (top) based on their DNA (DAPI) staining and the manual counting of their MUC2$^{+}$ cells. Scale bar, 100 µm (**M**) Fraction of MUC2$^{+}$ cells in mature colonic organoids quantified by 3D imaging with single-cell segmentation (12.81%, see Methods), by scRNA-seq (4.19% through combining all clusters expressing MUC2$^{+}$), and by FACS of fixed and MUC2-stained mature organoids.

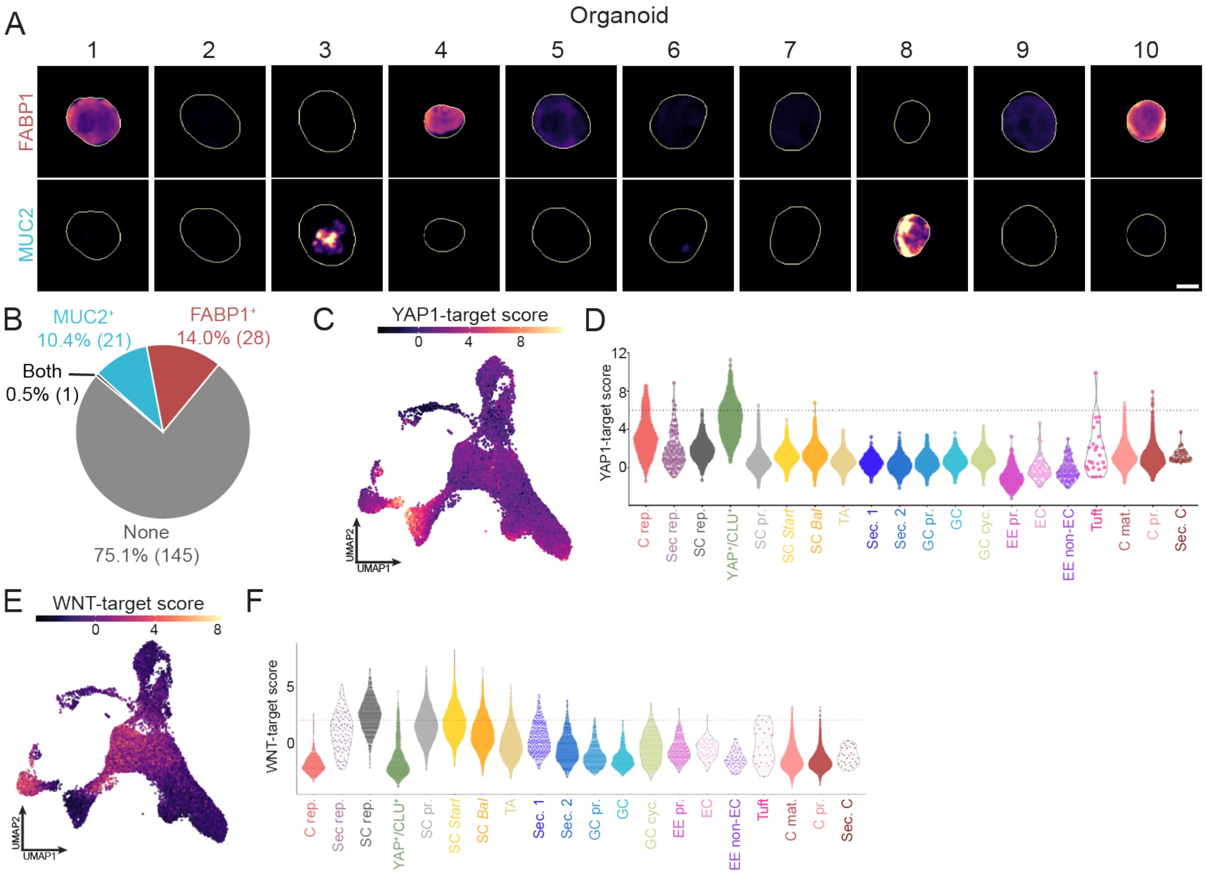


**Figure S5 | Organoid regeneration is driven by dedifferentiation trajectories.** ***Related to Figure 5*** (**A**) Images of ten randomly picked organoids of the first pseudotime bin from the imaging-based time course experiments (Figure 2) stained for FABP1 (upper row) and MUC2 (lower row) simultaneously. Images are maximum intensity projections (MIPs) of z-stacks acquired on a spinning-disk confocal system. Scale bar, 10 µm. (**B**) Quantification of organoids stained for FABP1 and MUC2 from the first pseudotime bin of the imaging-based time course experiments (Figure 2). Organoids were defined as $MUC2^{+}$ or $FABP1^{+}$ if their mean intensity reached a set threshold. If both thresholds are reached, organoids are defined as Both, while they were classified as None if neither threshold was reached. (**C**) UMAP of the transcriptomic landscape after cell-cycle correction and sketching (see Methods) color-coded by the YAP1-target gene score according to recent manuscripts[7,8]. (**D**) Violin plots depicting the YAP1-target score across transcriptomic clusters. (**E**) UMAP of the transcriptomic landscape after cell-cycle correction and sketching (see Methods) color-coded by the WNT-target gene score according to a recent manuscript[54]. (**F**) Violin plots depicting the WNT-target score across transcriptomic clusters.

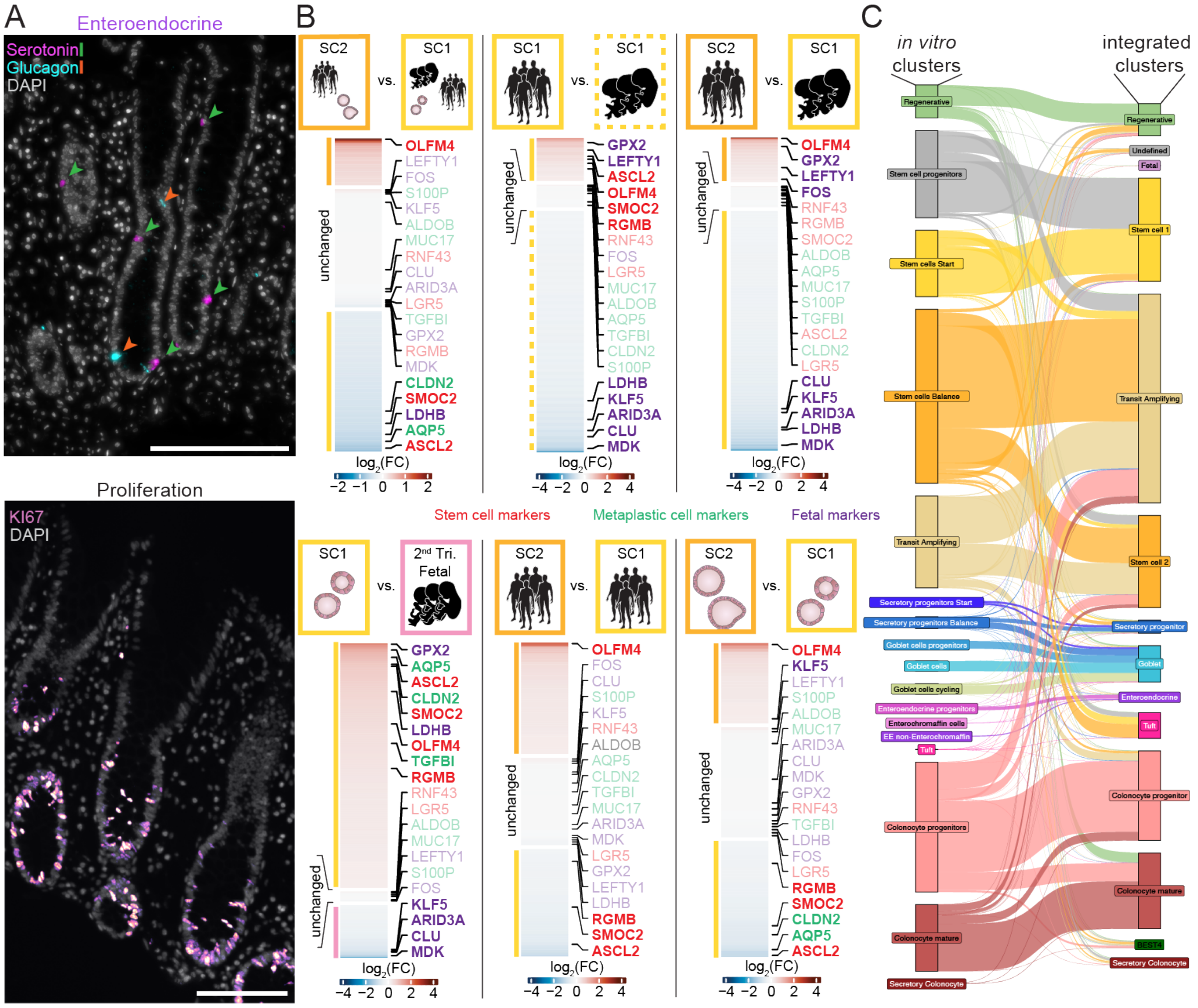


**Figure S6 | Colonic organoids resemble cell types of the *in vivo* tissue. *Related to Figure 6*** (**A**) Representative images of primary staining markers used in this study for enteroendocrine cells (Glucagon (orange) and Serotonin (green), top) and proliferation (KI67, bottom) in colonic tissue slices from biopsies of healthy individuals (see Methods). Images are maximum intensity projections (MIPs) of z-stacks acquired on a widefield microscope. Scale bar, 100 µm. (**B**) Differential gene expression analysis between stem cell clusters of the embedded scRNA-seq data from three *in vivo* studies and colonic organoids from this *in vitro* study. Shown is the comparison between the complete SC1 and SC2 set (left), the SC1 state between healthy adults and samples of the first trimester (second from left), the SC2 state of healthy adults and SC1 of first trimester samples (second from right) and SC1 of *in vitro* organoids and the Fetal cluster, consisting largely of cells from the second trimester of human fetal development (right). Genes are color-coded based on their reported association (stem cell genes: red, metaplastic markers: green, fetal markers: purple). (**C**) Sanky plot to showcase the connection between cells of the colonic *in vitro* scRNA-seq data set annotated in the stand-alone analysis (Figure 4, left side) and the embedded analysis (Figure 6, right side).

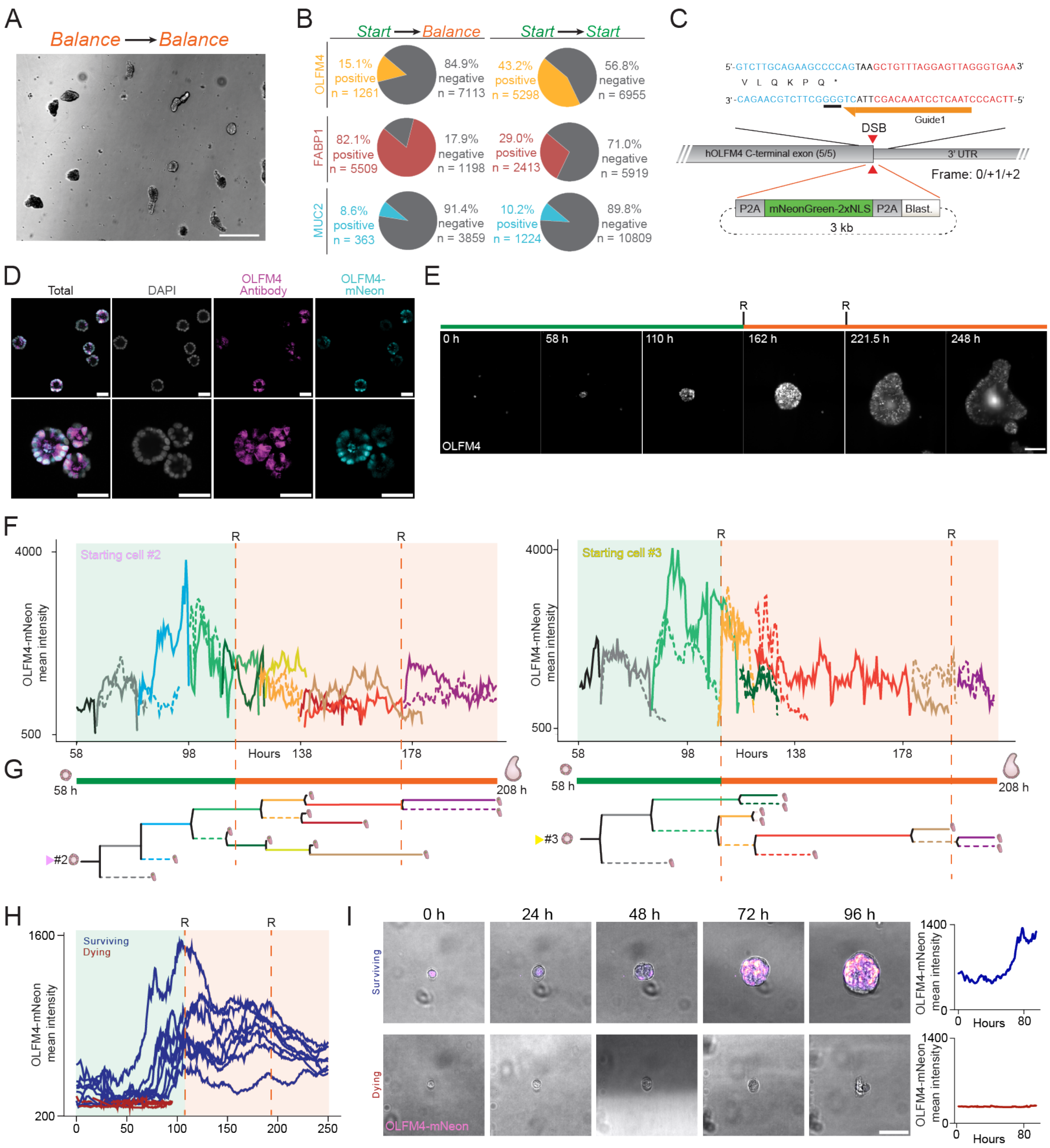


**Figure S7 | An initial first trimester-like stem cell state enhances intestinal regeneration. Related to Figure 7.** (**A**) Representative brightfield image of organoids grown purely in Balance medium. Scale bars, 100 μm. Related to Figure 7G. Due to the viability, no further features were analyzed. (**B**) Percentage as quantified by FACS after fixation and staining of mature organoids grown in the control condition (left) or purely in *Start* medium. Shown is the fraction of cells positive for OLFM4 (top row), FABP1 (middle row), or MUC2 (bottom row) among all analyzed cells. (**C**) Approach to genetically engineer an OLFM4-P2A-mNeon-NLS human colon reporter line (see Methods). (**D**) Representative images of the OLFM4-P2A-mNeon-NLS reporter line validation by comparing immunostaining against OLFM4 (magenta) with observed OLFM4-mNeon expression. Images are acquired on a spinning-disk confocal system. Scale bars, 50 μm. (**E**) Snapshots of an OLFM4-P2A-mNeon-NLS organoid imaged on a light sheet microscope for 250 hours with 30 min sampling intervals (Methods). The upper green bar indicates timeframe in *Start* medium, orange bar incubation in *Balance* medium. R indicates timepoints of medium refreshments. Images are maximum intensity projections (MIPs) of z-stacks. Scale bar, 100 μm. (**F-G**) Quantification of cell-level OLFM4-mNeon intensity along the organoid maturation trajectory on a light sheet microscope between 58 and 208 hours after seeding of single cells. Shown is the lineage of starting cell 2 out of 3 (left) and starting cell 3 out of 3 (right), related to Figures 7J-7L. Green background indicates timeframe in *Start* medium, orange background time in *Balance* medium. R and dashed lines indicate timepoints of medium refreshments. (**F**) Cell-lineage tree for highlighted cells in Figure 7J, starting cell 2 out of 3 left; starting cell 3 out of 3 right. Tracks are colored and stylized based on the cell id. Tracks which were lost are cells with expression levels of OLFM4 below the detection threshold, or cells with unclear movement of the cell. (**H**) Quantification of organoid-level OLFM4-mNeon intensity along the organoid maturation trajectory color-coded by the fate of the organoid (surviving: blue, dying: red) on a light sheet microscope over 250 hours. The tracks of all individual organoids are shown. Green background indicates timeframe in *Start* medium, orange background time in *Balance* medium. R and dashed lines indicate timepoints of medium refreshments. (**I**) Snapshots of the first 96 hours after single cell seeding of two OLFM4-P2A-mNeon-NLS organoids imaged on a light sheet microscope. Upper row shows an organoid which survives until the experimental endpoint at 250 h, while the lower row shows an organoid which died at 91 h. Line plots (right) show the mean OLFM4-mNeon expression of the exemplary organoid during the 96 h timeframe. Images are maximum intensity projections (MIPs) of z-stacks overlayed onto a brightfield image. Scale bar, 50 μm.

### Supplementary videos

**Video S1 | Human colon organoid maturation.** A human colonic nuclear reporter line (H2B-iRFP-670) imaged on a light sheet microscope (LS1, Viventis Microscopy) from 24 h after single cell seeding to 260 h in 15 min intervals. Shown is a MIP of a z-stack with 2 μm slicing-distance spanning the whole organoid.

**Video S2 | Human colon organoid maturation collage.** Collage of a human colonic nuclear reporter line (H2B-iRFP-670) imaged on a light sheet microscope (LS1, Viventis Microscopy) from 0 h after single cell seeding to 250 h in 30 min intervals. Shown is a MIP of a z-stack with 2 μm slicing-distance spanning the whole organoid.

**Video S3 | OLFM4 live dynamics of human colon organoid maturation.** Shown is a human colonic OLFM4 reporter line (OLFM4-P2A-mNeon-2xNLS) imaged on a light sheet microscope (LS1, Viventis Microscopy) from 0 h after single cell seeding to 200 h in 30 min intervals. Shown are MIPs of z-stacks with 2 μm slicing-distance spanning the whole organoid.

**Video S4 | Tracing of single cells in an OLFM4 human colon organoid reporter line.** Shown is a human colonic OLFM4 reporter line (OLFM4-P2A-mNeon-2xNLS) imaged on a light sheet microscope (LS1, Viventis Microscopy) from 58 h after single cell seeding to 208 h in 30 min intervals. Shown are MIPs of z-stacks with 2 μm slicing-distance spanning the whole organoid. Traces of three OLFM4+ starting cells color-coded by the age of the track (blue: oldest, red: most recent) are depicted. Related to Figures 7H, 7I, S7E, and S7F.

**Video S5 | OLFM4 live dynamics of human colon organoid maturation.** Shown is a human colonic OLFM4 reporter line (OLFM4-P2A-mNeon-2xNLS) which died while being imaged on a light sheet microscope (LS1, Viventis Microscopy). Movie from 0 h after single cell seeding until death at 96h in 30 min intervals. Shown are MIPs of z-stacks with 2 μm slicing-distance spanning the whole organoid. Lookup table histogram thresholds 10x lower compared to Video S3.

### Supplementary Notes

FACS strategy for OLFM4-mNeongreen-2xNLS positive cells

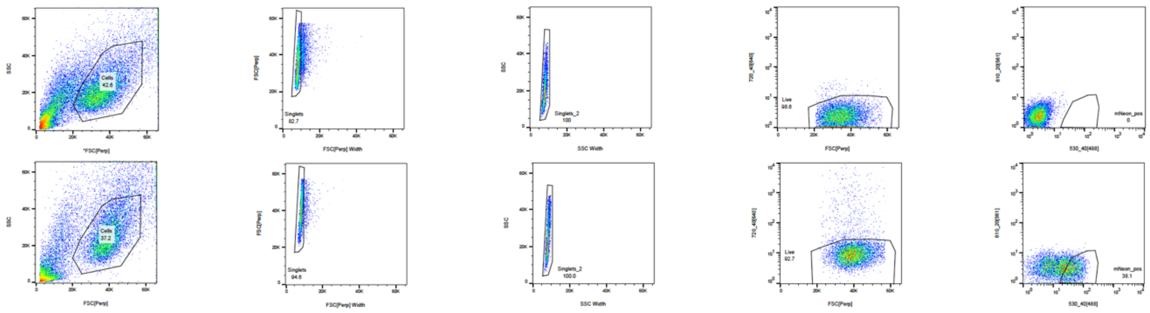

FACS strategy for pLV-H2B-iRFP positive cells

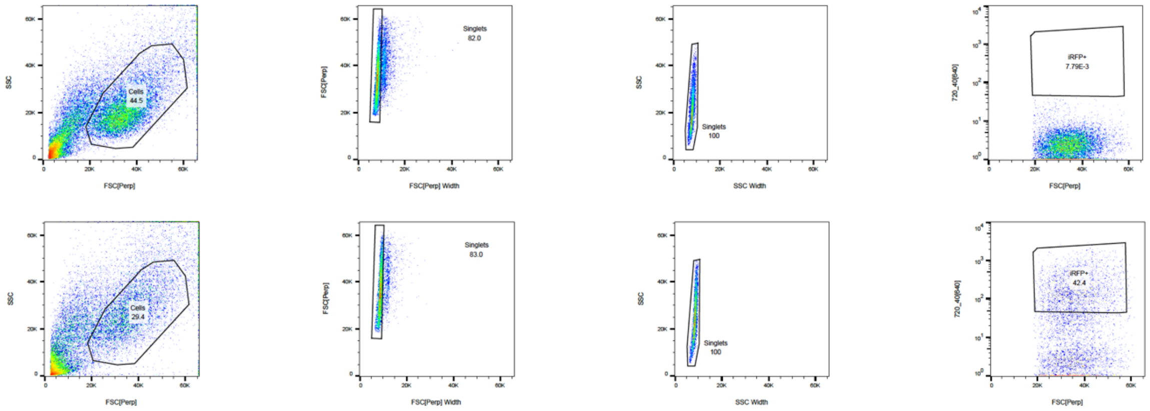

**Supplementary Note 1 | FACS strategy.** FACS plots to sort for OLFM4-mNeon-2xNLS positive (top) and H2B-iRFP positive (bottom) cells. Top row for both sorts show the control organoid line (without genetic engineering).

### Supplementary Tables

**Supplementary Table 1 | OLFM4 gRNAs.**

| | |
|---|---|
| **OLFM4_fwd** | CACCGAACTCCTAAACAGCTTACTG |
| **OLFM4_rev** | AAACCAGTAAGCTGTTTAGGAGTTC |
